# Evidence for trace gas metabolism and widespread antibiotic synthesis in an abiotically-driven, Antarctic soil ecosystem

**DOI:** 10.1101/2025.09.16.675928

**Authors:** A.R. Thompson, B.J. Adams, I.D. Hogg, S. Yooseph

## Abstract

The McMurdo Dry Valleys (MDVs) of Antarctica offer microbial ecologists a uniquely pristine, low-biodiversity model system for understanding fundamental ecological phenomena, the impact of a warming climate on ecosystem functioning, community structure and composition, and the dynamics of adaptation. Despite the scientific value of this system, we still know little about the functional ecology of its biota, especially the bacteria. Here, we analyzed the bacterial taxonomic and functional diversity of 18 shotgun metagenomes using the VEBA metagenome processing pipeline. We recovered 701 medium-to-high quality MAGs (≥ 50% completeness and contamination < 10%) and 201 high-quality MAGs (≥ 80% completeness and < 10% contamination), almost 50% more than found in similar sites previously. We found that: 1) community composition shifts along environmental gradients correlated with soil moisture, elevation, and distance to the coast; 2) many MDV bacteria are capable of performing trace gas metabolism; 3) genes associated with antibiotic-mediated competitive interactions (e.g., antibiotic biosynthesis and antibiotic resistance genes) are widespread; and 4) MDV bacteria employ survival strategies common to bacteria in similarly extreme environments. This study provides novel insight into microbial survival strategies in extreme environments and lays the groundwork for a more comprehensive understanding of the autecology of MDV bacteria.

## INTRODUCTION

The McMurdo Dry Valleys of Antarctica (MDVs) offer microbial ecologists an ideal model system for understanding fundamental ecological phenomena (Xue *et al*., 2024) and the effects of climate change on a microbial soil ecosystem (Fountain *et al*., 2014; Gooseff *et al*., 2017; Andriuzzi *et al*., 2018). The system is useful because of its reduced biotic diversity, its sensitivity to climate change, and the decades-long research focus by numerous research teams, including the United States National Science Foundation’s McMurdo Dry Valley Long-Term Ecological Research program (MCM LTER). Because of the harsh climate, there are no vascular plants, only three species of arthropods (Adams *et al*., 2006), and most soils (> 95%) host only a single metazoan (the nematode *Scottnema lindsayae*) or none at all (Virginia and Wall, 1999; Courtright *et al*., 2001; Bamforth *et al*., 2005). This is a system dominated by microbiota (Bamforth *et al*., 2005; Adams *et al*., 2006; Cary *et al*., 2010; Lee *et al*., 2012; Thompson *et al*., 2020). Soils are exceptionally arid (<5% soil moisture) (Burkins *et al*., 2001), saline, and oligotrophic, and as a polar desert the region is sensitive to climate change and likely to experience dramatic ecological changes sooner than many other ecosystems (Fountain *et al*., 2014; Gooseff *et al*., 2017). Moreover, research in these valleys has been ongoing for three decades and continues today with long-term experiments and data archives spanning more than 25 years (Gooseff *et al*., 2017; Andriuzzi *et al*., 2018). Researchers have investigated the diversity, distribution, taxonomy, and function of metazoans (Smith *et al*., 2012; Collins and Hogg, 2016; Shaw *et al*., 2018), bacteria (Cary *et al*., 2010; Sokol *et al*., 2013; Buelow *et al*., 2016), fungi (Fell *et al*., 2006; Gokul *et al*., 2013), protists (Bamforth *et al*., 2005; Thompson *et al*., 2020), archaea (Magalhães *et al*., 2014; Richter *et al*., 2014), and viruses (Zablocki *et al*., 2014; Wei *et al*., 2015; Adriaenssens *et al*., 2017; Bezuidt *et al*., 2020) as well as regional- and local-scale edaphic, climatic, geological, and hydrological parameters (Doran *et al*., 2002, 2010; Barrett *et al*., 2006, 2006; Simmons *et al*., 2009; Gooseff *et al*., 2017; Andriuzzi *et al*., 2018) .

Thus, MDV soils are a prime candidate for use as a comprehensive model of a soil ecosystem. However, key knowledge gaps remain regarding the functioning of these communities, limiting the full potential of this system as a model (Adams *et al*., 2006; Thompson *et al*., 2020, 2021; Xue *et al*., 2024). Perhaps most significantly, the genomic diversity and functional capabilities of its bacterial communities remain poorly understood.

Bacteria perform critical services to an ecosystem via nutrient cycling and community structuring processes (Sikorski, 2015). These services are mediated by the diversity of a community’s ecological functioning and life history strategies, many of which are influenced by bacterial phylogenetic and taxonomic diversity. Thus, characterizing bacterial taxonomic and functional diversity can inform aspects of ecosystem functioning. It is well established that MDV soil communities exist along a climatic gradient that somewhat correlates to the distance from the coast and elevation and exerts some control over their composition and structure (Marchant and Head, 2007; Dragone *et al*., 2022; Mashamaite *et al*., 2023). What is less clear is how this gradient impacts low-level taxonomic diversity (genus and species) and functional attributes (Van Horn *et al*., 2013; Dragone *et al*., 2022), such as nutrient cycling and survival strategies.

Understanding this relationship is important as previous research has predicted that as the MDVs warm due to climate change, the more arid soil communities will be replaced by taxa common in current moist soils (Andriuzzi *et al*., 2018), which may threaten the potentially unique taxonomic and functional diversity of arid MDV communities.

How the majority of MDV soil communities outside algal mats, moss beds, and lithic environments is sourcing their carbon (whether from atmospheric trace gases, windblown mat material, or legacy sources) is debated (Ortiz *et al*., 2021). Recent work has shown that trace gas metabolism—the incorporation of CO_2_, CO, and H2 via non-photosynthetic means—is widespread in soils globally (Bay *et al*., 2021) as well as in Antarctic sites similar to those of the MDV (Ji *et al*., 2017; Ortiz *et al*., 2021). However, to our knowledge, no study has confirmed this for sites that have been the focus of comprehensive characterization. Nitrogen is also limited in the MDVs (Monteiro *et al*., 2020), with previous studies unable to recover nitrogen fixing microbes outside wetted margins of ponds and streams (Cary *et al*., 2010).

It is unclear how much MDV soil communities are structured by biotic and abiotic variables (Caruso *et al*., 2019; Lee *et al*., 2019; Thompson *et al*., 2021). Multi-tiered trophic levels exist in MDV soils (Bamforth *et al*., 2005; Shaw *et al*., 2018; Thompson *et al*., 2021) and studies have identified antibiotic-associated genes from various surveys (Fierer *et al*., 2012; Wei *et al*., 2015; Núñez-Montero and Barrientos, 2018). Whether bacteria are competing directly (via antagonistic interactions) or indirectly (via extremotolerance survival strategies) is still an unanswered question.

To corroborate prior estimates of MDV bacteria diversity and assess the role of MDV soil bacteria in ecosystem functioning across a climate gradient, we analyzed a dataset of 18 soil metagenomes previously used to characterize the taxonomic diversity and functional roles of soil eukaryotes in the MDVs (Thompson *et al*., 2020, 2021). Here, we: (1) evaluate bacterial community structure and function across climatic gradients; (2) examine the genetic potential for bacterial trace gas metabolism and nitrogen and sulfur metabolism; (3) estimate the potential for antibiotic-mediated antagonistic interactions; and (4) identify tolerance strategies that enable survival in the harsh Antarctic environment. To our knowledge, this is the first genome-resolved metagenomic study of bacteria from these widely surveyed sites in the MDVs

## METHODS

### Soil sample collection, DNA extraction, and sequencing

Metagenomes were generated and analyzed previously for their eukaryotic component (Thompson *et al*., 2020, 2021). Here we investigate the prokaryotic component of the metagenomes using genome mapping and the VEBA software for genome annotation (Espinoza and Dupont, 2022; Espinoza *et al*., 2024). Full details on sampling and sequencing are provided in Thompson *et al*. (2020, 2021) and (Beet *et al*., 2016). Briefly, 87 soil samples were taken from 18 sites comprising a representative range of landscape types and features, latitudes, and environmental gradients (Fig 1, Table S3). Most samples consisted of undeveloped mineral soils though several possessed varying degrees of vegetation (i.e., Hjorth Hill, moss; Cliff Nunatak, algae, and Canada Stream, biocrust). Samples were collected between 2014 and 2017 using sterilized scoops and sterile Whirl-Pak® bags and stored for transport and long-term storage at -20°C. Subsamples were used to evaluate moisture content, pH, electrical conductivity, total N, total C, total P, NO_3_-N. Texture and climatic zone was assigned to each site using available literature (Fountain *et al*., 2014; Thompson *et al*., 2020). Due to financial constraints, samples were pooled prior to DNA extraction, which was performed using the DNeasy PowerSoil Kit (Qiagen) with protocol optimized for low biomass soils (see Thompson *et al*., [2020] for details). Libraries were prepared with NEBNext Ultra II DNA Prep Kit (New England Biolabs) and sequenced on an Illumina HiSeq 2500 at a read length of 2 x 250 bp and a total insert length of 500 bp (Thompson *et al*., 2020). See supplemental materials for run stats (Table S1).

**Figure 1.**
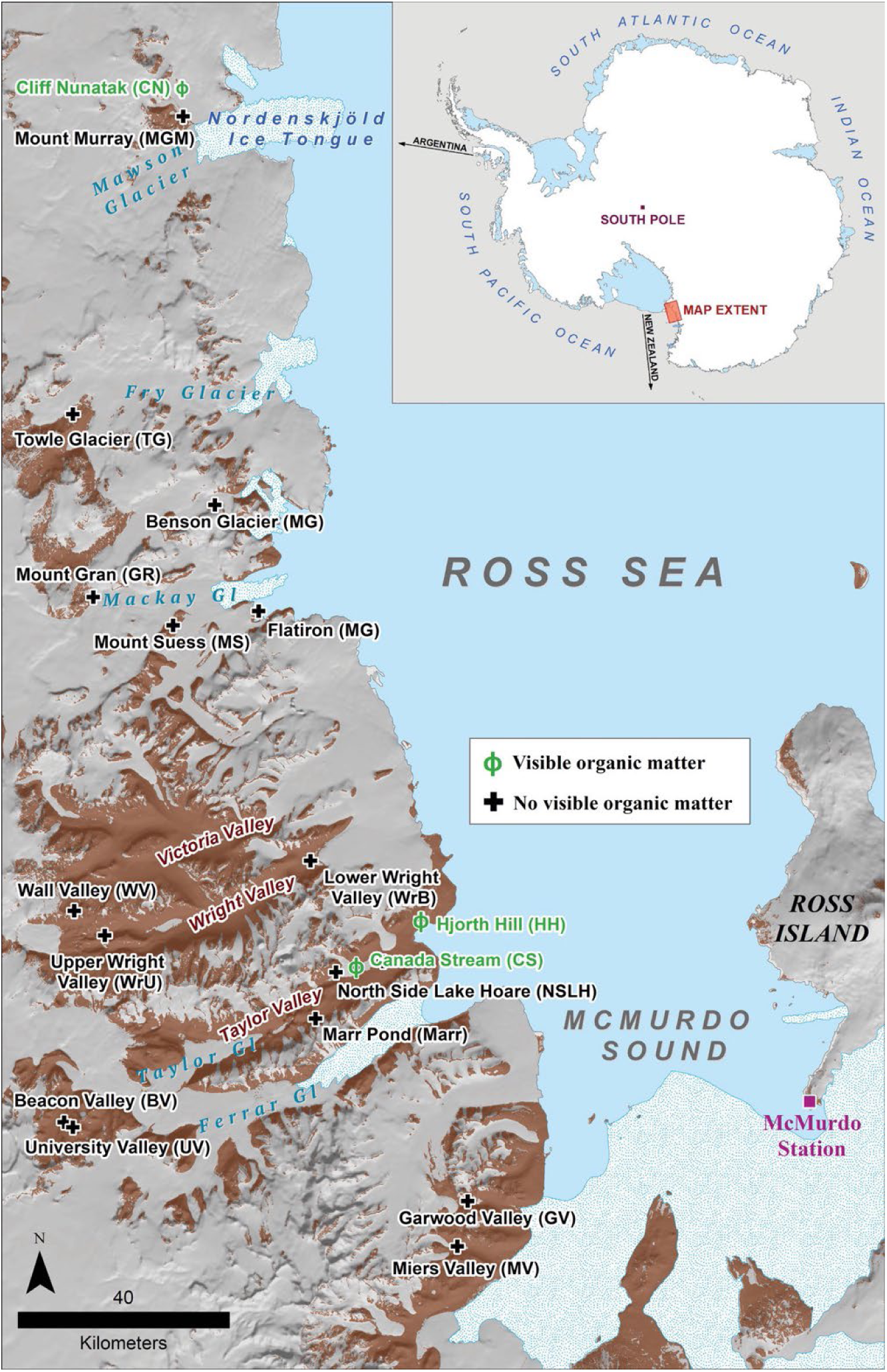
– Map of study area (Reproduced with permission from Thompson et al. 2020). The McMurdo Dry Valleys are located at roughly S 77° E 162° in Southern Victoria Land and open towards the Ross Sea to the East. Samples represent a variety of MDV soil habitats and landscape features that exist in the MDVs. Samples in green (Φ) possessed visible organic matter (e.g. moss and algae); samples in black were coarse to fine mineral soils with varying degrees of visible moisture. Figure by Mike Cloutier, Polar Geospatial Center.

### Genome Database Construction and Mapping

To assess the relative abundances of microbial genomes in our metagenomes, we mapped quality trimmed sequence reads from each sample to a collection of reference genomes (Jones *et al*., 2015; Loftus *et al*., 2021). Reads were trimmed using Trimmomatic with the settings LEADING 2, TRAILING 2, SLIDINGWINDOW 4:15 (default), MINLEN 30 (Bolger *et al*., 2014; Thompson *et al*., 2020). For this analysis, we built a custom database using genomes sourced from publicly available databases, then mapped our reads to these genomes using Bowtie2 (Langmead and Salzberg, 2012). The read mapping information was analyzed using a probabilistic framework based on a mixture model to estimate the relative copy number of each reference genome in a sample (Xia *et al*., 2011; Loftus *et al*., 2021). To build the reference database, we used refSeq (O’Leary *et al*., 2016), JGI GOLD (Mukherjee *et al*., 2025), and a google scholar literature search for Antarctic prokaryote genomes using combinations of the following characteristics: “soil”, “desert”, “cold desert”, “antarctic”, “polar”, “psychrotolerant”, and “assemble”. We included genomes from maritime Antarctic islands and Antarctic marine samples because there is overlapping diversity between continental, maritime, and marine zones (Cary *et al*., 2010; Varliero *et al*., 2024). We used NCBI’s batch download, filtered for complete genomes and those with annotations, excluded anomalous assemblies, those from metagenomes, and from unverified source organisms. Our resulting database (after removing duplicate genomes) contained 79,686 total genomes (78,082 refseq Bacteria genomes, 1,066 refseq Archaea genomes, 526 genomes from the JGI Environment search, and 12 from the Antarctic literature search).

### Metagenome processing

Raw, demultiplexed sequences were processed using the Viral Eukaryotic Bacterial Archaea (VEBA) open-source pipeline, an end-to-end metagenomics software that can preprocess, assemble, cluster, classify, and produce preliminary statistics on metagenomes from all domains of life and viruses (Espinoza and Dupont, 2022; Espinoza *et al*., 2024). The pipeline was run with default settings for all modules. Briefly, sequences were trimmed using *fastp* (Chen *et al*., 2018) and assembled using *metaSPAdes* (Nurk *et al*., 2017). Unassembled reads were next mapped back to the assemblies using *Bowtie2* (Langmead and Salzberg, 2012), and then *SeqKit* (Shen *et al*., 2016) was used to assess assembly statistics. Binning was performed by integrating multiple group-specific software to optimize functionality and accuracy. For prokaryotes, *CoverM* (Aroney *et al*., 2024) was used to estimate coverage; *MaxBin2*, *MetaBat2*, and *CONCOCT* were used for binning (Alneberg *et al*., 2014; Wu *et al*., 2016; Aramaki *et al*., 2019); *DAS Tool* was used for dereplication (Sieber *et al*., 2018); *Tiara* was used for filtering out non-prokaryotic sequences (Karlicki *et al*., 2022); *CheckM* was used for quality assessment and filtering (Parks *et al*., 2015); and *Prodigal* and GTDB-Tk were used for gene calling and genome classification, respectively (Hyatt *et al*., 2010; Chaumeil *et al*., 2020). VEBA removed poor quality MAGs according to its defaults: completeness ≤ 50% or contamination > 10 (Espinoza and Dupont, 2022). MAGs were subsequently clustered into species-level clusters (SLCs) at a 95% threshold (default) using *FastANI* (Jain *et al*., 2018). To annotate MAG proteins, *Diamond* (Buchfink *et al*., 2015) was used to align protein sequences against the NCBI non-redundant protein database (default) while protein domains were identified using HMMER (Mistry *et al*., 2013) and the *Pfam* database (Mistry *et al*., 2021) and KEGG orthology was assessed using *KOFAMSCAN* (Aramaki T *et al*.). Finally, reads were mapped back to assemblies to estimate gene feature counts using the index building and mapping features in *Bowtie2*. The pipeline also processed contigs identified as eukaryotic or viral, though these data were not analyzed for this study.

### Analyses

All analyses were conducted in R version 4.4.1 (R Core Team, 2024) and plots were generated using ggplot2 version 3.5.1 (Wickham, 2016) unless specified otherwise. To visualize MAG taxonomic diversity and assembly quality, we created a Sankey diagram (Fig. 2) with the ggalluvial R package, version 0.12.5 (Brunson and Read, 2023). Assembly quality stats and taxonomic classifications for the Sankey diagram were sourced from the CheckM and taxonomic_classification files produced by VEBA (Espinoza *et al*., 2024), which were combined then filtered for the top 25 taxa at each taxonomic level.

**Figure 2.**
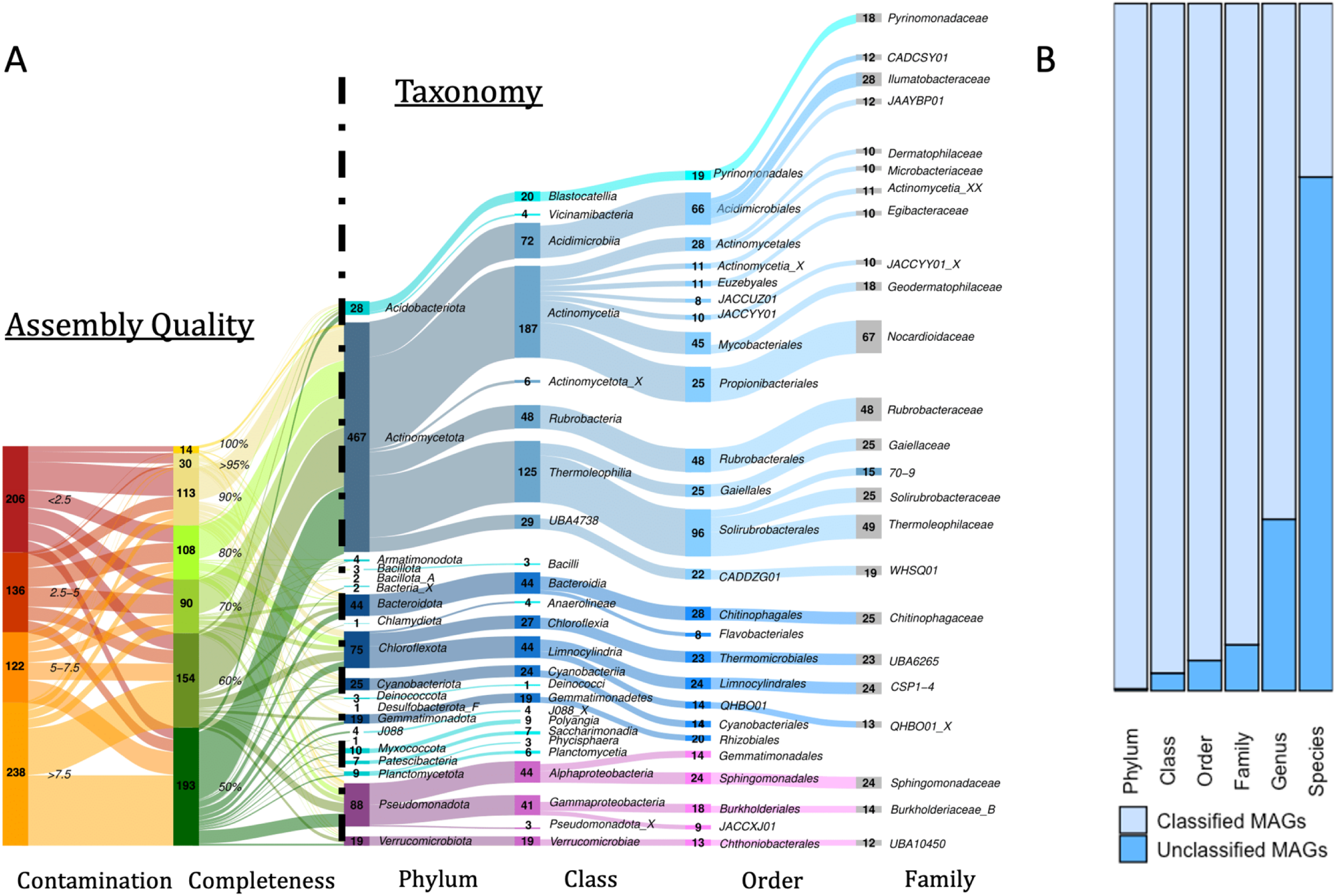
– MAG taxonomy and assembly stats. A) Sankey diagram of MAG taxonomic diversity coupled to assembly completeness (yellows and greens) and contamination level (reds and oranges). Only the top 20 taxa are shown for each level, and genera and species were excluded from the visualization for clarity. Completeness percentages and contamination levels are shown in italics to the right of each category’s counts, respectively. Completeness percentages represent ranges; for example, ’50%’ includes all assemblies from 50% (inclusive) to 60%. (exclusive). B) Proportion of classified to unclassified MAGs for each taxonomic level. Unclassified MAGs are defined as lacking a classification in the VEBA output and are displayed as “taxon_X” in the Sankey diagram.

To visualize community dissimilarity, PCAs of CLR-transformed species-level clustered MAG and ORF counts were made using prcomp() function in base R (Fig. 3). To visualize the relative abundance of taxa, heatmaps of CLR-transformed (Nearing *et al*., 2022) species-level clustered MAG counts were made using pheatmap version 1.0.12 (Kolde, 2018) (Fig. 4).

**Figure 3.**
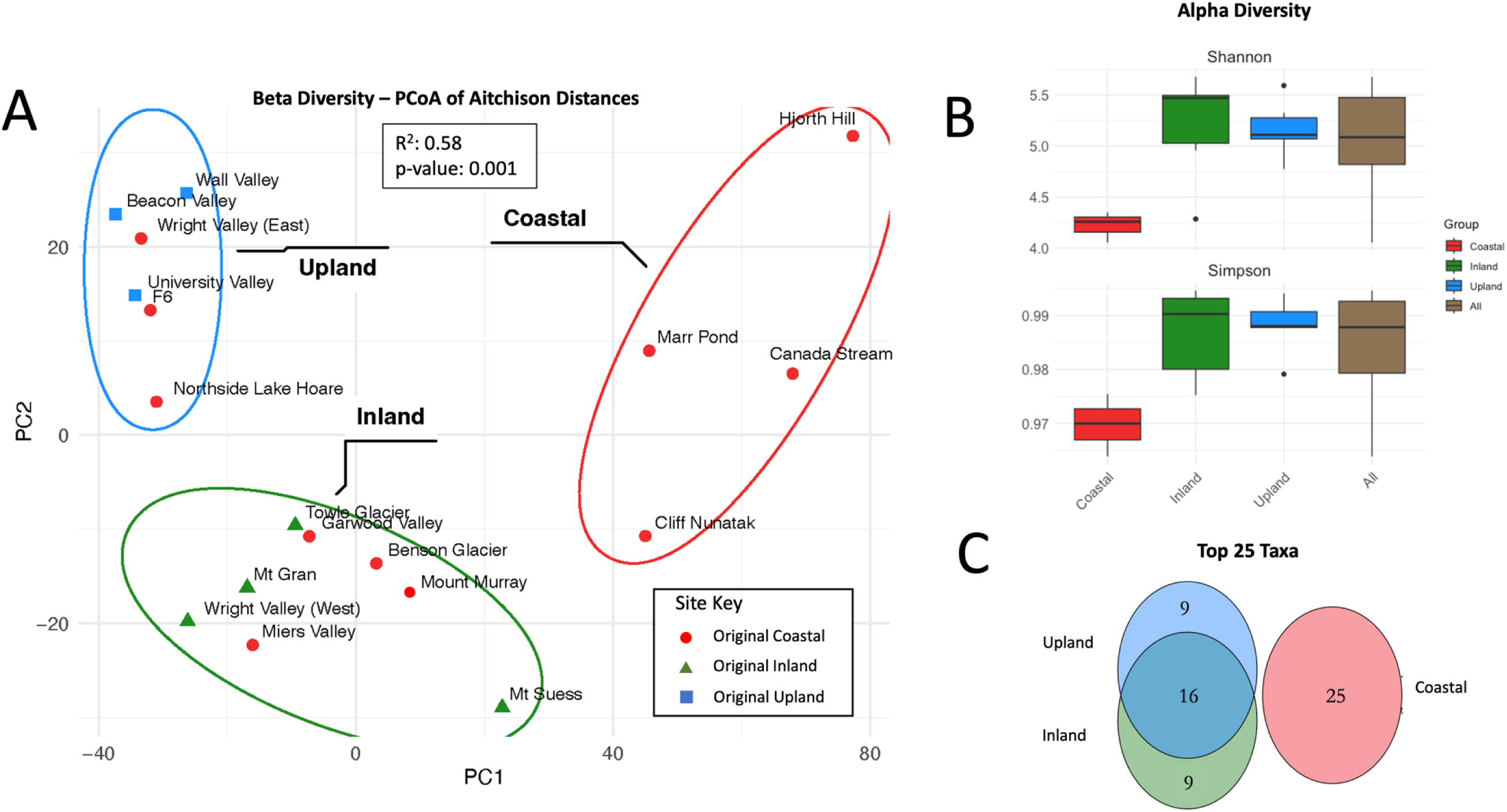
– Diversity metrics and taxonomic overlap. A) Beta diversity: PCA of site dissimilarity based on clr-transformed MAGs (species-level; SLCs), colored by climatic zones: coastal – red; inland – green, upland – blue. Site icons designate previous climactic designations: coastal – red circle; inland – green triangle; upland – blue square. Ellipses around sites represent statistically significant site groupings. B) Alpha diversity by zone and total (Shannon and Simpson’s indices) using relativized counts. C) Venn diagram showing overlap between top 25 most abundant taxa by climatic zone.

**Figure 4.**
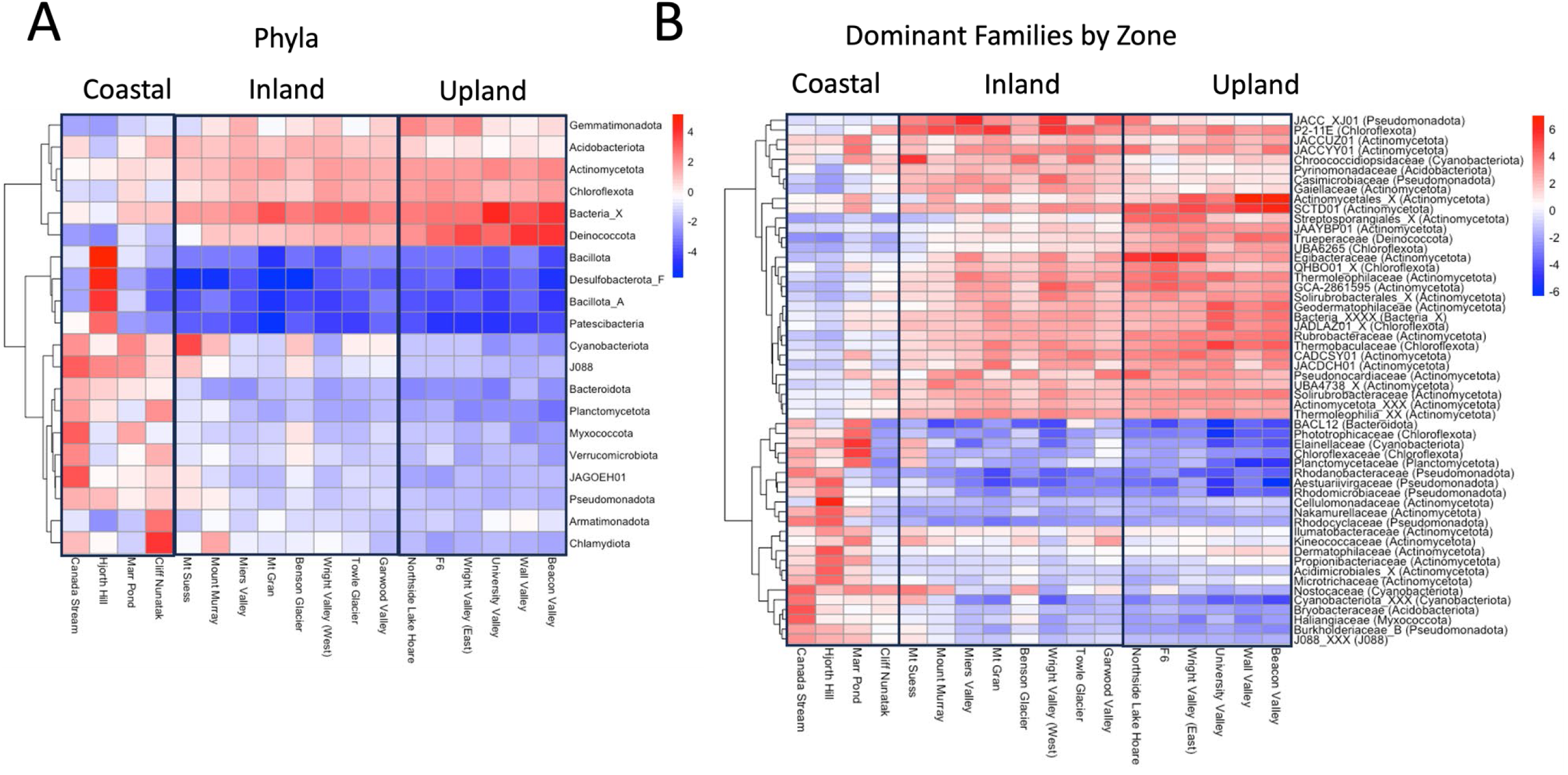
– Heatmap of clustered MAGs (species-level, SLCs) counts against sites at phylum (A) and family levels (B). Family levels represent top 50 most relatively abundant taxa in each climatic zone and taxon labels are family names followed by phylum in parentheses. Sites are grouped by climate groupings as estimated in Figure 3 (left to right: coastal, inland, upland). Made using CLR-transformed MAG counts data.

KEGG pathway completion ratios produced by VEBA were used to produce Figures 5 through 8. Mean completion values for each pathway was calculated by averaging completion ratios for each listed pathway across sites (either all sites for Figures 5 and 6 or sites in each climate zone for Figures 7 and 8). For climate zone specific Figures 7 and 8, the top 25 most abundant taxa from each zone were used (Fig. S5). KEGG pathways were restricted to those involved in potentially ecologically significant behaviors including virulence factors, extremotolerance, competition, or nutrient cycling, using the literature.

**Figure 5.**
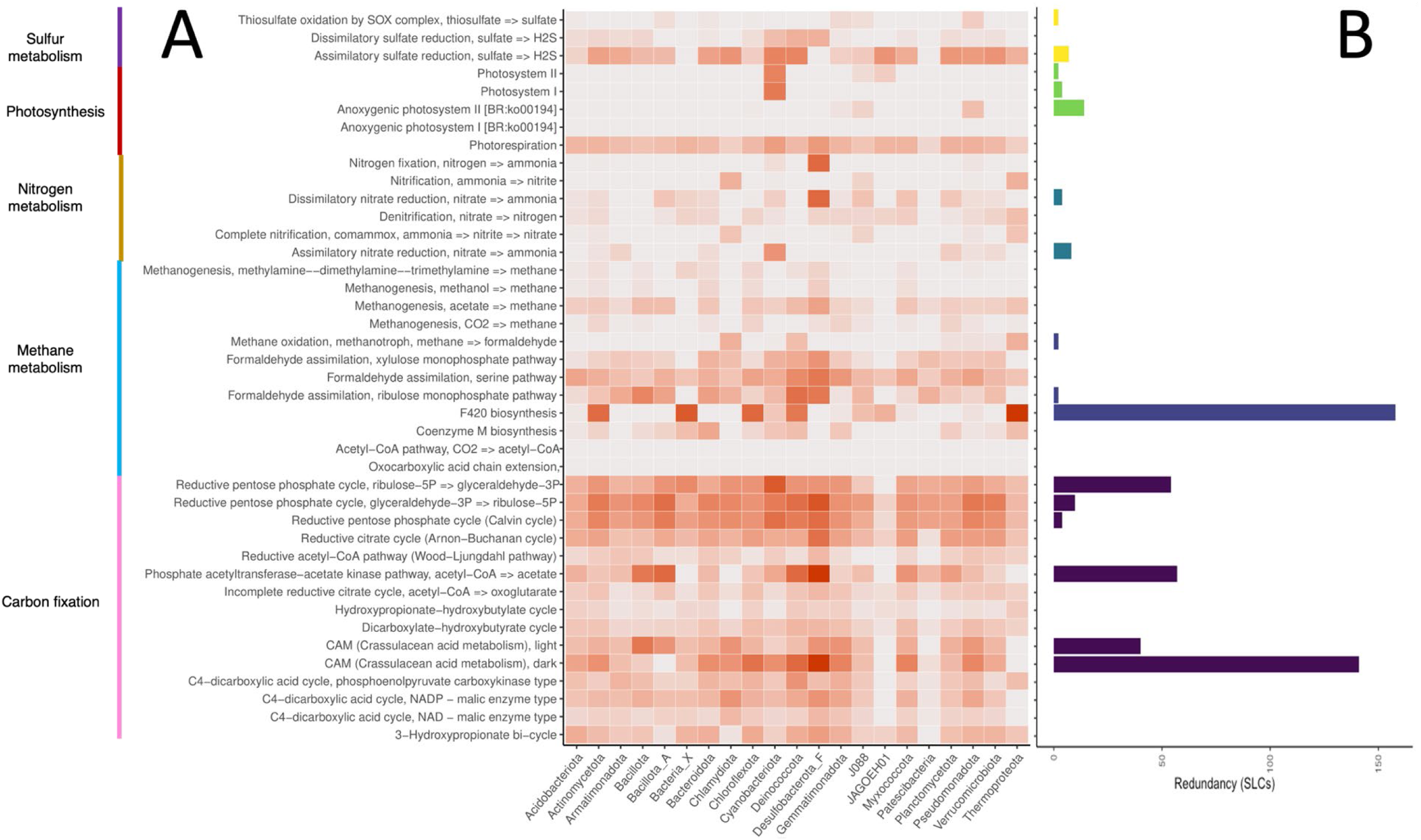
– Completion and redundancy of KEGG modules associated with metabolism of key nutrients. A) Mean completion of each pathway module by bacterial phylum. The darker the orange, the closer completion ratio is to 100%. B) Number of SLCs with 100% complete KEGG gene pathways by module, a proxy for functional redundancy. Blank rows indicate no SLC possessed that pathway to 100% completion in any site. X-axis represents phyla and taxonomic count, respectively. Y-axis represents KEGG pathways and modules.

**Figure 6.**
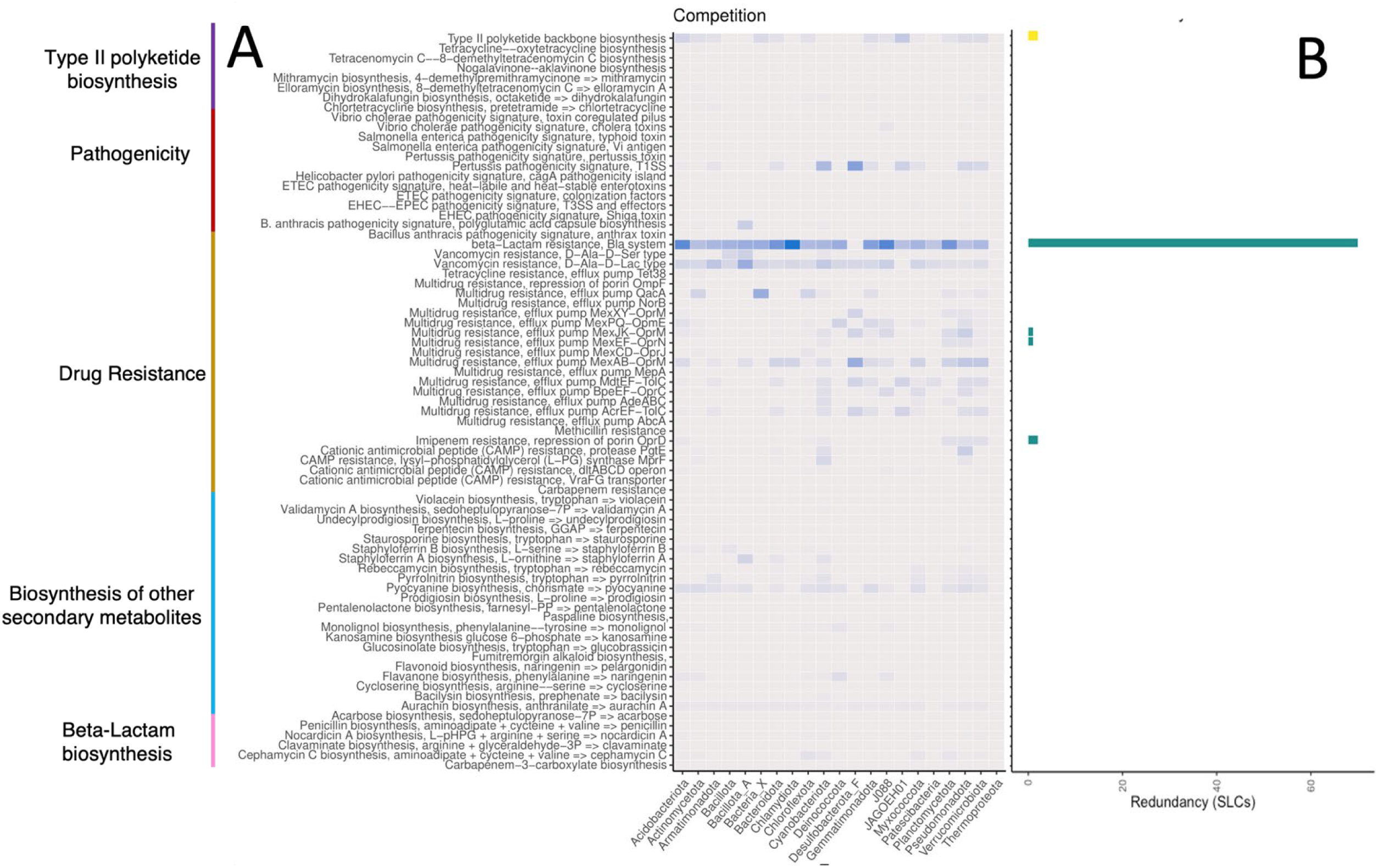
– Completion and redundancy of KEGG modules associated with inter-species competition. A) Mean completion of each pathway module by bacterial phylum. The darker the blue, the closer completion ratio is to 100%. B) Number of SLCs with 100% complete KEGG gene pathways by module, a proxy for functional redundancy. Blank rows indicate no SLC possessed that pathway to 100% completion in any site. X-axis represents phyla and taxonomic count, respectively. Y-axis represents KEGG pathways and modules.

**Figure 7.**
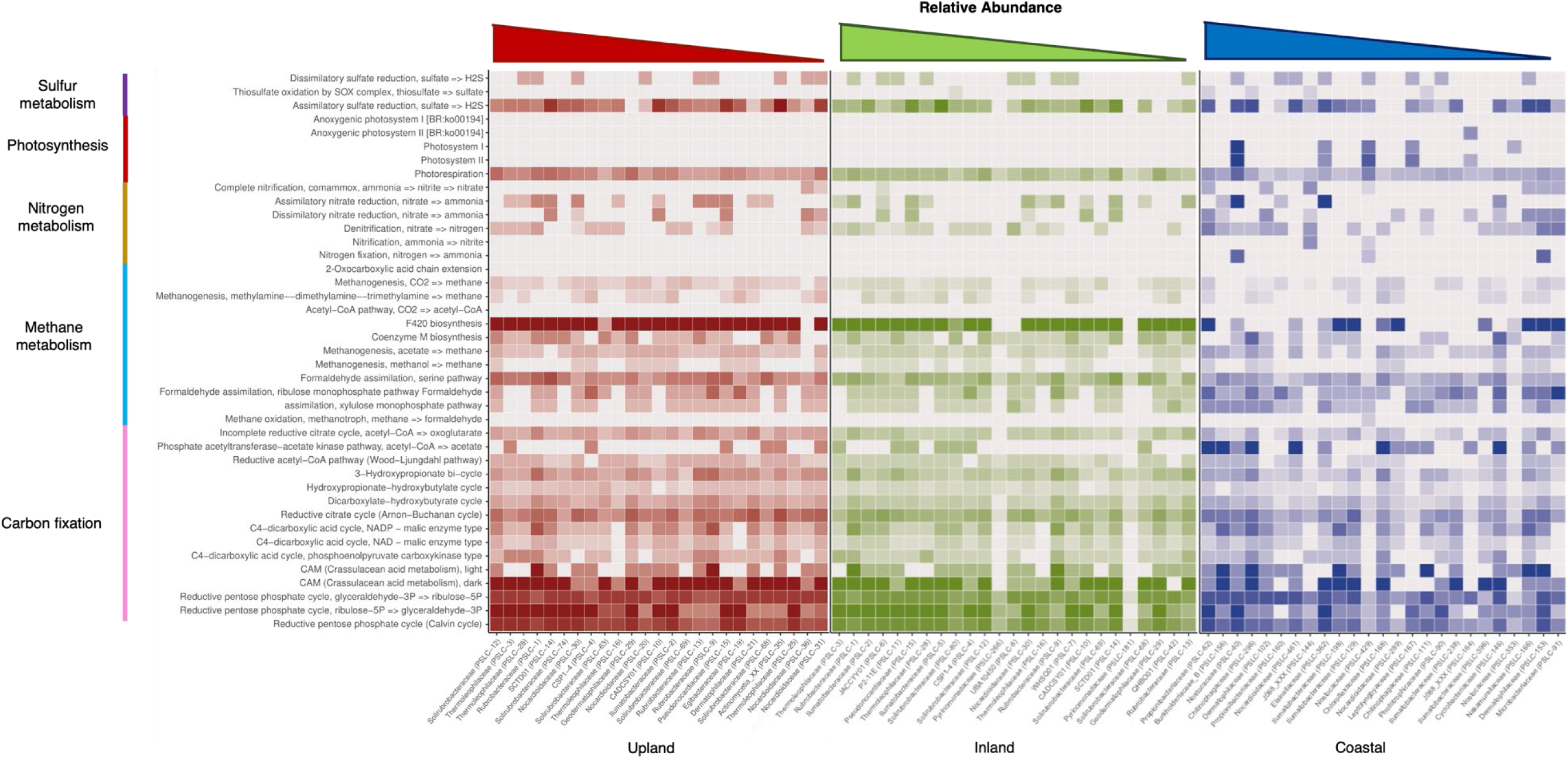
– Metabolic functional differences between climatic zones (for top 25 abundant taxa). Functional differences are estimated using completion ratios of KEGG modules displayed as heatmaps. Climatic zones are as follows: inland (red), upland (green), and coastal (blue). Taxa (x-axis) represent the most relatively abundant taxa by zone (based on clr-transformed SLC counts) with relative abundance increasing towards the left side of each graph. Taxa are listed with family followed by the unique SLC taxon id assigned by VEBA (e.g., PSLC-1) for convenience in understanding their functional ecology and for comparing dominant taxa composition across zones. Y-axis shows KEGG pathways (far left) and select module names for each pathway (near left).

**Figure 8.**
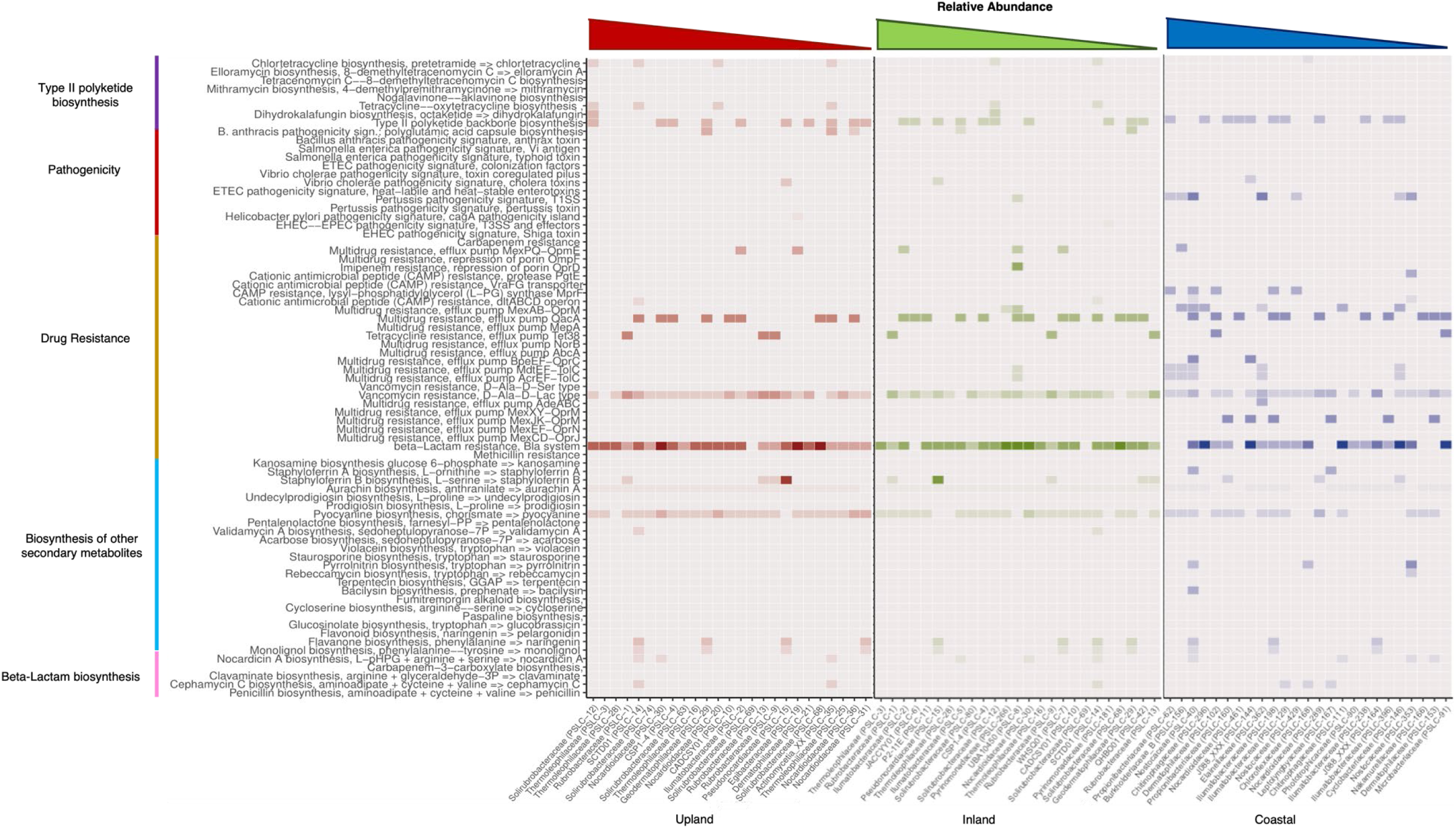
– Competitive functional differences between climatic zones (for top 25 abundant taxa). Functional differences are estimated using completion ratios of KEGG modules displayed as heatmaps. Climatic zones are as follows: inland (red), upland (green), and coastal (blue). Taxa (x-axis) represent the most relatively abundant taxa by zone (based on clr-transformed SLC counts) with relative abundance increasing towards the left side of each graph. Taxa are listed with family followed by the unique SLC taxon ID assigned by VEBA (e.g., PSLC-1) for convenience in understanding their functional ecology and for comparing dominant taxa composition across zones. Y-axis shows KEGG pathways (far left) and select module names for each pathway (near left).

A literature search was performed to identify keywords of genes and phenotypes involved in desiccation tolerance, cold shock response and tolerance, osmotic stress tolerance, radiation tolerance, antibiotic resistance, antibiotic synthesis, virulence factors (Chandra and Kumar, 2017; Cycoń *et al*., 2019), and the six known pathways for fixing carbon (e.g., Calvin-Benson cycle and Arnon-Buchanan cycle; see Berg [2011]). The list of keywords resulting from the search (Table S2) were queried against our gene annotation tables to estimate phenotype frequency in our MAGs.

## RESULTS

### Taxonomic Diversity

Using default settings, VEBA recovered 701 medium-to-high quality bacterial MAGs (completeness > 50% and contamination < 10%) and 201 high quality bacterial MAGs (completeness > 80% and contamination < 10%). The 701 medium-to-high quality MAGs clustered to 485 species-level MAG clusters (SLCs) in 20 phyla, 38 classes, 82 orders, and 140 families. Most MAGs (75%) were unidentifiable at the species level. However, 75% corresponded to previously identified genera and 93% to previously identified families. There was a negative relationship between genome contamination and completeness (-0.109, R^2^ = 0.2779, p-value = <2e-16; Fig. S1) and the average genome size was 3,336 genes and the average GC content was 64.58 %. Additionally, an average of 3.1% of sequences and 12.8% bps remained unbinned after processing (Table S1).

Eight phyla (Actinomycetota, Pseudomonodota [previously Proteobacteria], Chloroflexota, Bacteroidota, Acidobacteriota, Cyanobacteriota, Gemmatimonadota, and Verrucomicrobiota) of the twenty total phyla accounted for 93% of the MAGs. The majority (55%) of MAGs belonged to the phylum Actinomycetota, while Pseudomonadota (12%) and Chloroflexota (10%) together comprised nearly half (49%) of the remainder. Five taxonomic identifications were resolved to the species level: *Pseudomonas E antarctica*, *P. E gergormendelli*, *Carnobacterium A pleistocenium, C. A alterfunditum*, and *Flavobacterium fryxellicola* (data not shown). We also recovered a single Archaeal MAG, an unclassified *Nitrosocosmicus* (phylum Thermoproteota, order Nitrosphaerales) that was 83.65% complete and had 2.85% contamination (data not shown).

Only a small fraction of the reads mapped to the genomes in our custom database (Fig. S3). Reads mapped better to the JGI GOLD sequences (∼ 40%) than to refseq (∼ 28%) and surprisingly poorly to genomes from the Antarctic literature (∼ 4%). The vast majority of reads did not map with any existing sequences, while an average of ∼23,000 (0.12% of the total) reads mapped once and 14,000 (0.08%) mapped more than once (Fig. S3).

### Community structure by climatic zone

All MAGs occurred in all sites, but community structure depended on site climatic zone (Fig. 3). Site communities clustered into three groups (PERMANOVA; p-value: 0.001, R^2^: 0.58) that loosely corresponded to the site’s previously determined climatic zones (Marchant and Head, 2007; Thompson *et al*., 2020). Over half (60%) of sites determined as belonging to the coastal zone (near the coast, low elevation) clustered instead with inland or upland sites (Fig. 3A). For the purposes of this paper, we reassigned zones based on Fig. 3A. Diversity in coastal sites was significantly different than from inland and upland sites according to both the Shannon and Simpson diversity indices (Kruskal-Wallis Nonparametric test; p-value: 0.38 and 0.39, respectively; Pairwise tests: p-values: 0.36 and 0.38, respectively). Clusters were dominated by distinct taxa, though there was overlap between the taxonomic profiles of the inland and upland site while there was none between the coastal cluster communities and the inland and upland communities.

At the phylum level, community structure differed substantially between the coastal cluster and the upland and inland clusters, but was highly consistent between the inland and upland sites. An unclassified phylum (Bacteria_X), Deinococcota, Gemmatimonadota, Acidobacteriota, Actinomycetota, and Chloroflexota were the most abundant phyla in the inland and upland zone communities, except for Marr Pond, Cliff Nunatak, and Mt Seuss. Conversely, the phyla Bacillota, Desulfobacteriota_F, Bacillota_A, and Patescibacteria had the lowest relative abundance at all sites except Hjorth Hill, where they were the most abundant. The coastal sites were dominated by Cyanobacteria, J088, Myxococcota, JAGOEH01, Pseudomonadota, and Chlamydiota (Fig. 4A).

At lower taxonomic levels (i.e., family), the structure of the inland and upland clusters became distinct (but remained relatively homogenous internally) and sites in the coastal cluster showed significant differences to one another (Fig. 4B). Two unnamed groups, P2-11 E (Chloroflexota) and JACCXJ01 were consistently the most abundant families in the inland zone, except for Chroococcidiopsidaeceae (Cyanobacteriota) at Mount Suess. Egibacteriaceae, an unnamed group SCDTD01, and an unclassified Actinomycetales (all Actinomycetota) were highly abundant across the upland cluster (Fig. 4B).

When considering the 25 most abundant taxa per zone, the upland and inland zones shared 16 of their top 25 taxa. However, all of the top 25 taxa from the coastal zone were unique to that zone (Fig. 3C). The structure of the top 25 taxa in each zone was characterized by a change from Actinomycetota to non-Actinomycetota bacteria; Actinomycetota were 24 of the top 25 taxa in the upland zone, 19 in the inland zone, and 13 of the coastal sites. However, the 13 Actinomycetota from the coastal sites were mostly different families (Illumatobacteraceae, Propionibacteriaceae, and Dermatophilaceae, and Nocardioidaceae) than those found in the upland and inland zones (Solirubrobacteraceae, Thermoleophilaceae, Nocardioidaceae, and Rubrobacteraceae). Nocardioidaceae and Illumatobacteraceae were the only families to occur in the top 25 taxa in all three zones, though there was no overlap between the top 25 coastal taxa and the taxa from the other zones at the species level (Fig. 6). The non-Actinomycetota in the coastal zone consisted of phyla Cyanobacteria (families Nostocacaeae, Leptolyngbyaceae, and Elainellaceae), Bacteroidota, Chloroflexota, J088, and Pseudomonodota.

### Functional Ecology of MDV soil bacteria communities

Overall, few taxa possessed the complete set of genes for key KEGG modules in these categories, although many possessed partially complete pathways for those modules (Fig. 5B). Modules contributing to inter-species competition (e.g., secondary metabolites, virulence factors, antibiotic biosynthesis, and antibiotic resistance) were likewise rarely complete and also infrequently partially complete (Fig. 6B).

### Nutrient metabolism

For carbon metabolism, most taxa possessed some genes involved in each CO_2_ metabolism KEGG pathways (Calvin cycle [99.4%], Arnon-Buchanan cycle [100%], Wood-Ljungdahl pathway [71%], the Hydroxypropionate-hydroxybutylate cycle [79%], 3-Hydroxypropionate bicycle [98%], and the Dicarboxylate-hydroxybutyrate cycle [98%]).

However, average completion was low across phyla (Fig. 5A; less than 25% completion for all pathways except the Arnon-Buchanan [41%] and Calvin [56%] cycles) and none had a 100% complete gene set for any of these pathways except four taxa for the Calvin cycle (two alphaproteobacterial [Bradyrhizobium, Sphingomonas] and two gammaproteobacteria [Dokdonella, Rhodoferax]). An additional 39 taxa had 91% complete KEGG Calvin cycle genesets. Of the other carbon fixation cycles, the Arnon-Buchanan cycle pathway was most complete across taxa, with 148 SLCs possessing >= 50% of the pathway’s genes, 28 >= 70%, and 3 >= 80% of the pathway genes (a Pseudoxanthomonas [Xanthomonodaceae, Gammaproteobacteria]; and unidentified taxa in the Dermatophilaceae [Actinomycetota] and Opitutaceae [Verrucomicrobiota]).

We recovered hits against at least one gene considered potentially diagnostic of each cycle (Berg, 2011) in all cycles except the dicarboxylate-hydroxybutyrate cycle (Fig. 9A). Three cycles had key genes in a significant proportion of taxa: 178 taxa had phosphoribulokinase (Calvin cycle), 335 taxa had propionyl-CoA carboxylase (Hydroxypropionate [Fuchs-Holo] bi-cycle), and 315 taxa had fumarate reductase (Arnon-Buchanan cycle), though only two had taxa which possessed all enzymes associated with the cycle (the Wood-Ljungdahl pathway [17] and the 3-hydroxypropionate/4-hydroxybutyrate (HP/HB) cycle [14]). Additionally, a number of genes coding for CO metabolism (formate and CO dehydrogenase) and H2 gas oxidation ([Ni-Fe]- and [FeFe]-hydrogenase) occurred in roughly half and one quarter of the SLCs, respectively.

**Figure 9.**
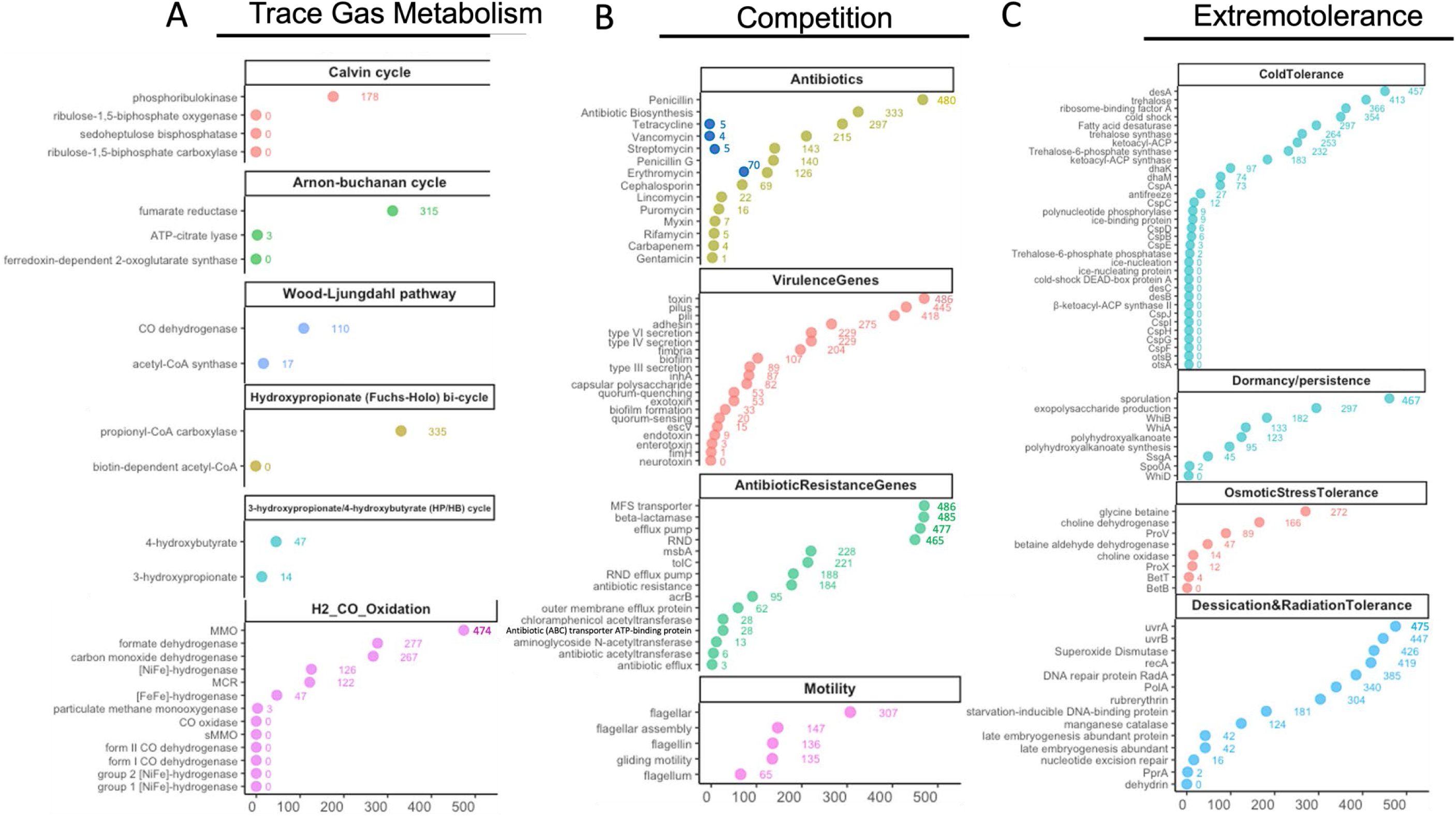
– Phenotypic Keyword Hits by Taxa. Phenotypes are organized by category, with gene keywords (e.g., antibiotic acetyltransferase) on the y-axis and taxa counts on x-axis. Counts (x-axis) represent unique taxa with at least 1 hit for any given phenotype/gene. Keywords with 0 hits were excluded from figure; (A) trace gas metabolism, (B) competition, and (C) extremotolerance.

For nitrogen metabolism, only members of desulfobacteriota possessed near complete KEGG pathways for nitrogen fixation, while some portions of the KEGG nitrification pathway occurred in the bacterial phyla Chlamydiota and J088 and the archaeal phylum Thermoproteota (Fig. 5). Nitrate reduction and denitrification were more common across phyla but 100% completion of the pathway in KEGG was rare (Fig. 5). Assimilatory nitrate reduction was more widespread among the top 25 taxa of the inland and upland zones than of the coastal (12 and 12 versus 5, respectively). In contrast, dissimilatory nitrate reduction was more common in coastal site taxa than in the inland and upland zones (11 versus 6 and 5, respectively; Fig. 7). Nitrogen fixation and nitrification via ammonia pathway genes were absent from the top 25 taxa in the inland and upland zones, while nitrification via commamox was more common in the coastal sites than nitrification via ammonia, and rare outside the coastal zone (Fig. 7).

### Competitive interactions

Few KEGG modules associated with competitive interactions were complete or even partially complete (Fig. 6). Only beta-lactam resistance was broadly present and mostly complete across phyla and dominant taxa in each zone (Fig. 8), with more than 60 taxa possessing 100% complete KEGG gene suite for the pathway (Fig. 6B). Vancomycin resistance (D-Ala-D-Lac type) and pyocyanine biosynthesis were just as widely distributed as beta-lactam resistance, although much less complete (no taxon had a fully complete gene suite for these pathways; Fig. 6). Type II polyketide backbone biosynthesis and multidrug resistance efflux pump QacA were sporadically present in phyla from all zones, and were likewise incomplete in all taxa (Figs 6B, 8). Overall, there was more competition-associated functional diversity in coastal-dominant taxa than there was in the upland or inland zones with genes from nine modules uniquely in the coastal zone compared to four in the upland/inland zone (Fig. 8).

Our gene annotation file queries provided additional insight. We recovered hits on 13 of 77 antibiotic keywords (Table S2) in our annotation files, seven of which occurred in more than 100 MAGs (Fig. 9). Of those that did occur across taxa, all but one (erythromycin) consisted of mostly hits pointing to genes involved in resistance to those compounds. Erythromycin returned more than half of its gene hits containing the string “erythromycin biosynthesis”. To estimate the number of distinct taxa that possessed biosynthesis genes for these widely occurring antibiotics, we added additional queries to our list combining the antibiotic’s name and “biosynthesis”, then added the results to Figure 9B as dark blue circles.

Beyond antibiotics, genes associated with toxin production and pili were nearly ubiquitous across taxa (486 and 445, respectively), while adherence and type IV and IV secretion systems were less common but still present in roughly half of the MAGs. General antibiotic resistance phenotypes (major family system transporters, efflux pumps, and RND) were widespread, while phenotypes that targeted specific antibiotics (e.g., various acetyltransferases) were rarer. Additionally, at least 28% of MAGs possessed genes associated with flagella, and an equal number with gliding motility. Slightly fewer (∼20%) possessed genes containing the keyword biofilm.

### Extremotolerance genes

We queried four categories of extremotolerance genes (cold tolerance, dormancy/persistence, osmotic stress tolerance, and desiccation & radiation tolerance) against our annotation files to ascertain which strategies MDV bacteria were using to survive in the MDVs. Cold tolerance strategies including fatty acid desaturase (and the accompanying gene desA), trehalose, and cold shock associated proteins, were found in a majority (54%, 95%, 85%, 61%, respectively) of MAGs, while sporulation and exopolysaccharide production were common (96% and 61%, respectively) dormancy strategies. Over half of MDV taxa also possessed glycine betaine and choline dehydrogenase genes, both involved in the production and export of the osmoprotectant glycine betaine. Genes and phenotypes typically associated with DNA damage repair (uvrAB, recA), and reactive oxygen species scavenging (superoxide dismutase) were also nearly ubiquitous strategies for managing the effects of desiccation and irradiation.

## DISCUSSION

### Taxonomic Diversity (MAGs diversity and assembly completeness)

Our results reveal a diverse and previously under-characterized bacterial community in MDV soils, with 701 medium-to-high quality MAGs corresponding to an estimated 484 unique species-level genomes spanning distinct environmental gradients. This is 55% more MAGs than previously recovered in a study of similar sites (Ortiz *et al*., 2021), suggesting that microbial diversity in polar desert soils exceeds that previously recognized. Though this discrepancy in MAG count between studies could be an artefact from the differences in sequencing depth, sampling breadth, and binning approaches (Vosloo *et al*., 2021), our study used VEBA, a pipeline which uses a novel iterative prokaryote binning procedure that tends to recover additional MAGs from sites previously analyzed with other approaches (Espinoza and Dupont, 2022). Actinomycetota dominate this ecosystem, consistent with previous observations of Antarctic sites (Cary *et al*., 2010; Ji *et al*., 2017). Two MAGs belonged to phyla that GTDB-tk was unable to classify. Both were from an environmentally extreme high elevation valley that has been used for astrobiology research (Goordial *et al*., 2016) and both were near the lower threshold for medium quality assemblies (∼51% completeness and > 6% contamination), suggesting the presence of novel yet rare extremophile lineages in the more extreme MDV soils. We also recovered the nitrogen-oxidizing archaeon *Nitrosocosmicus* from the order Nitrososphaerales and phylum Thermoproteota (Alves *et al*., 2019), a less abundant yet consistently present archaeon in this region’s soils (Ortiz *et al*., 2021; Dragone *et al*., 2022).

Intriguingly, one of the most abundant families in the upland zone (SCTD01) has also been reported from fire-combusted soils in temperate climates (Nelson *et al*., 2022).

Few MAGs were resolved to genus or species level, few KEGG pathways for key metabolic and cell maintenance processes were complete, and few of our reads mapped well to genomic sequences in standard databases. This apparent genetic novelty could be attributed to the fact that two thirds (62%) of our MAGs were less than 80% complete and thus lacked genomic information that these species actually possess. However, 17% (84/481) of the species level clusters (SLCs) had two or more MAGs assigned to them, increasing the likelihood that we captured much more of the genome than the minimum of 50% across the MAGs for each of these species. Additionally, 24 of the 25 most abundant taxa had multiple MAGs per SLC, and their KEGG completion profiles (Figs 7 & 8) show a similar partial completion pattern to all the taxa (Figs 5 & 6). A high degree of genetic novelty would not be surprising as Antarctic soil communities are highly isolated both geographically and temporally (Convey *et al*., 2008, 2009), Antarctic bacteria are genetically distinct at fine taxonomic scales (Albanese *et al*., 2021), and standard databases represent environmental isolates poorly, particularly from less studied environments (McGee *et al*., 2019). Our results support the notion that further exploration of Antarctic soil bacterial genomes and phenotypes will likely yield previously unknown metabolic pathways and survival strategies (Rizzo *et al*., 2021; Waschulin *et al*., 2022). Such knowledge can provide insights into ecological genomics (Xue *et al*., 2024), applied microbiology (Mesbah, 2022), human health (Núñez-Montero and Barrientos, 2018), and astrobiology (Rawat *et al*., 2024).

### Community structure and functional ecology

MDV soils appear to be dominated by two general communities with distinct, albeit related, functional profiles, correlating to long-term moist areas (e.g., coastal sites) and long-term arid sites (i.e., inland and upland sites). The long-term arid sites are dominated by gram-positive Actinomycetota and possess a more homogenous functional profile, while the long-term moist sites possessed greater phylogenetic and functional diversity, but these patterns are likely to shift with climate change.

### Nutrient metabolism

We found widespread evidence for trace gas metabolism as a key survival strategy. A substantial number of MDV bacteria possess key genes for fixing atmospheric carbon non-photosynthetically via the Calvin and Arnon-Buchanan cycles, and the Hydroxypropionate (Fuchs-Holo) bi-cycle. While many taxa contained genes for multiple carbon fixation pathways, few possessed fully complete KEGG pathways, suggesting that mix-and-match metabolic flexibility may be common in these resource-limited soils (Ortiz *et al*., 2021). These results align with recent studies emphasizing the role of atmospheric trace gas oxidation in polar desert microbiomes (Ji *et al*., 2017; Ortiz *et al*., 2021) and soils globally (Bay *et al*., 2021) and confirms previous inferences (e.g., Ortiz *et al*., 2021) with higher taxonomic resolution.

We confirm that nitrogen fixation does not appear to occur outside moist sites (Cary *et al*., 2010), although taxa capable of performing assimilatory nitrate reduction are more common in arid soils than in moist sites.

### Competitive interactions and extremotolerance

Based on the genetic composition of our MAGs, we suggest that antagonistic interactions between MDV bacterial taxa are a critical factor in MDV community dynamics. Genes associated with antibiotic biosynthesis (primarily in the form of antibiotic biosynthesis monooxygenase) occurred in nearly 70% of taxa, suggesting that the capability for antibiotic production is widespread. Additionally, antibiotic resistance genes (for penicillin, vancomycin, and streptomycin especially) were common across taxa, though vancomycin and streptomycin production were not, suggesting that antibiotic resistance is critical to survival in MDV soil communities. While functional annotation suggests widespread resistance genes, the limited detection of complete biosynthetic pathways suggests novel compounds. Nonetheless, few taxa possessed genes associated with typical antibiotic compounds (Table S2), which suggests there may be novel antibiotics in Antarctic soil bacteria. This observation potentially conflicts with broader observations that abiotic factors typically outweigh biotic pressures in structuring Antarctic soil communities (Hogg *et al*., 2006; Lee *et al*., 2019; Lebre *et al*., 2023), but is qualitative rather than quantitative and requires additional investigation to contextualize it.

Our data also confirm that MDV bacteria invest heavily in genetic and physiological extreme tolerance survival mechanisms, suggesting that selection for taxa that can manage both biotic and abiotic pressures must be high. Many MAGs possessed genes predicted to be involved in managing low temperatures (fatty acid desaturase), long-term unfavorable conditions (sporulation), ice nucleation (glycine betaine, trehalose, and exopolysaccharide production), and DNA damage repair and ROS scavenging (uvrAB and superoxide dismutase). Few MAGs possessed the genes often associated with these cellular processes (e.g., otsAB for trehalose synthase, CSPA-J for the cold shock response) (Lim *et al*., 2019; Zhang and Gross, 2021; Scales *et al*., 2023), again suggesting that MDV bacteria are utilizing novel gene pathways for key survival phenotypes. However, this observation could also be the result of non-standard gene naming in databases, erroneous assemblies, or other limitations of predictive gene annotation using standard reference databases (Salzberg, 2019).

Finally, many taxa possessed genes for motility features, such as flagella and gliding motility, as well as genes with the keyword “biofilm”. However, these phenotypes were found in only a subset of MAGs, suggesting that MDV bacteria lifestyles are diverse and that the soils they inhabit are likely to contain several micro-niches (Vos *et al*., 2013; Larkin and Martiny, 2017; Baquero *et al*., 2021).

## CONCLUSIONS

In conclusion, we recovered 50% more medium- and high-quality MAGs than previously reported for similar Antarctic sites. We also report MAGs from sites in the same features as those studied as part of the MCM LTER, expanding the known genomic diversity and multifunctionality of these ecosystems. Our findings challenge the appropriateness of using previously designated climatic zones as proxies for microbial community structure, suggesting that localized abiotic and biotic factors may play a larger role in shaping microbial assemblages. However, we suggest that community composition remains distinct between moist, vegetated sites and arid, regolith-based sites. We provide strong evidence for diverse carbon fixation strategies in MDV taxa, extending beyond photosynthetically driven carbon fixation and spanning multiple climatic zones. Our data suggest that interspecies competition, particularly antibiotic-mediated interactions, may be a significant structuring force in microbial food web dynamics in these soils. Finally, our results support previous findings that Antarctic bacteria likely possess numerous novel genes and pathways involved in key nutrient metabolism pathways, extremotolerance, and antibiotic biosynthesis. These insights underscore the potential for unique microbial adaptations in extreme polar desert environments and highlight the need for further exploration of their functional roles in Antarctic ecosystems.

## ACKNOWLEDGEMENTS

This research was funded by the National Science Foundation (NSF) Grant numbers ANT 2133685 and OPP-2224760 to BJA and is a contribution to the McMurdo Dry Valleys Long Term Ecological Research (LTER) program. Antarctica New Zealand and the New Zealand Antarctic Research Institute (NZARI) provided logistics and financial support through Event K024 to IDH. This research was also funded by the NSF grant #DBI-2400009 awarded to Shibu Yooseph. Additional support was provided by the Monte L. Bean Life Science Museum, the Department of Biology, Brigham Young University and by the Kravis Department of Integrated Sciences, Claremont Mckenna College. Geospatial support for this work provided by the Polar Geospatial Center under NSF-OPP awards 1043681, 1559691, and 2129685.

## AUTHOR CONTRIBUTIONS

Contributed to conception and design: AT, SY. Contributed to analysis of data: AT. Contributed to interpretation of data: AT, BA, SY. Drafted and/or revised the article: AT, BA, SY, IH. Approved the submitted version for publication: AT, BA, SY, IH.

## DATA AVAILABILITY

Raw metagenomes are publicly available on NCBI under Project Title “WGS 18MDV 2018” All other data and R code is available on GitHub at: https://github.com/Andy-Thmpsn/18MDV_analysis_code/18MDV_Bacteria_MAGs.R

