## Supplemental Materials for "Evidence for trace gas metabolism and widespread antibiotic synthesis in an abiotically-driven, Antarctic soil ecosystem"

**Supplemental Figures**

Supp. Table 1 – Sequencing Stats

Supp. Table 2 – Phenotype Keywords

Supp. Table 3 – Sample Physicochemical & Climatic Zone Metadata

Supp. Table 4 – Alpha & Beta Diversity Stats

Figure S1 – MAG and SLC genome size (est. gene count)

Figure S2 – Metadata Overview

Figure S3 – Alignment Success & Protein divergence*

Figure S4 – KEGG Completion Ratio Boxplots

Figure S5 – Taxon relative abundance distribution

Supp. Table 1 – Sequencing, Assembly, and Binning Stats

| Sample Information | | Sequencing Stats | | | | Assembly (contigs) | | | | | | | Binning | |
|---|---|---|---|---|---|---|---|---|---|---|---|---|---|---|
| Sample Name | Samples Pooled | Soil Extracted (g) | DNA submitted (pg/ul) | R1 Reads (Mb) | Run Proportion | Sequences in assembly (M) | Total length of sequences (Gb) | minimum length | average length | maximum length | N50 | GC(%) | % unbinned (seqs) | % unbinned (bp) |
| Mount Gran * | 1 | 5.25 | 2047.5 | 7.5 | 2.74% | 1.6 | 870.9 | 56 | 531.9 | 86837 | 529 | 65.1 | 3.0% | 11.9% |
| Towle Glacier * | 3 | 5.25 | 2120.4 | 8.8 | 3.21% | 2.1 | 1093.9 | 55 | 513.5 | 43286 | 491 | 63.83 | 2.9% | 11.7% |
| Lower Wright Valley | 8 | 5.25 | 2044 | 11.5 | 4.18% | 1.6 | 836.9 | 55 | 522 | 242066 | 539 | 64.95 | 3.4% | 14.4% |
| Mount Murray | 3 | 5.25 | 2039.8 | 11.6 | 4.24% | 3.4 | 1675.8 | 55 | 486.2 | 539558 | 450 | 64.68 | 2.0% | 8.2% |
| East Side Lake Fryxell/F6 | 2 | 5.25 | 2112.6 | 12.8 | 4.68% | 1.6 | 857.8 | 55 | 521 | 74203 | 590 | 65.99 | 3.3% | 13.8% |
| Upper Wright Valley * | 8 | 7 | 2057.09 | 13.0 | 4.74% | 2.6 | 1381.9 | 56 | 535 | 131499 | 542 | 65.05 | 4.0% | 15.7% |
| Garwood Valley | 6 | 7 | 2209.9 | 13.2 | 4.82% | 3.2 | 1464.0 | 56 | 463.4 | 39961 | 452 | 64.01 | 2.0% | 8.9% |
| Wall Valley * | 8 | 7 | 2084.6 | 13.2 | 4.83% | 1.3 | 727.3 | 55 | 540.9 | 59786 | 623 | 63.69 | 4.8% | 20.2% |
| University Valley * | 7 | 10.5 | 1304.05 | 13.2 | 4.83% | 1.3 | 773.6 | 56 | 594.7 | 121249 | 698 | 66.35 | 4.4% | 18.6% |
| Marr Pond | 4 | 5.25 | 2081.52 | 13.3 | 4.84% | 2.9 | 1544.6 | 56 | 534.4 | 383571 | 512 | 60.86 | 2.4% | 8.8% |
| Mount Suess | 3 | 5.25 | 2179.2 | 15.5 | 5.66% | 4.0 | 2018.3 | 55 | 503.1 | 304172 | 467 | 61.78 | 2.5% | 9.8% |
| North Side Lake Hoare | 3 | 5.25 | 2025 | 16.0 | 5.83% | 2.4 | 1133.8 | 55 | 479.7 | 93731 | 548 | 65.1 | 3.0% | 13.9% |
| Benson Glacier/Flatiron | 3 | 3.5 | 2037 | 16.3 | 5.96% | 4.3 | 2122.2 | 55 | 497.3 | 165713 | 466 | 62.23 | 2.4% | 9.6% |
| Miers Valley | 6 | 7 | 2076.36 | 17.1 | 6.23% | 3.5 | 1770.9 | 56 | 503 | 130205 | 493 | 65.16 | 3.2% | 13.8% |
| Cliff Nunatak ϕ | 2 | 3.5 | 2073.9 | 21.8 | 7.95% | 4.2 | 2355.5 | 55 | 562.1 | 545376 | 547 | 61.23 | 3.3% | 12.2% |
| Canada Stream ϕ | 1 | 1.75 | 2100 | 21.8 | 7.96% | 5.8 | 2978.1 | 55 | 516.8 | 347067 | 482 | 58.61 | 2.1% | 7.7% |
| Hjorth Hill ϕ | 5 | 3.5 | 2043 | 21.9 | 8.01% | 2.9 | 1681.2 | 55 | 571.5 | 330481 | 551 | 60.15 | 2.6% | 12.1% |
| Beacon Valley * | 14 | 10.5 | 2175 | 25.4 | 9.28% | 2.1 | 1203.3 | 55 | 560.5 | 170084 | 671 | 64.73 | 4.6% | 19.2% |
| Mean | 5 | 6 | 2045 | 15.2 | 6% | 2.8 | 1471.7 | 55.3 | 524.3 | 211602.5 | 536 | 63.5 | 3.1% | 12.8% |
| Total | 87 | 103.25 | | 273.8 | 100% | 50.9 | 26489.9 | | | | | | 56% | |

Table S1 – Sequencing stats from Thompson, A. R., S. Geisen, and B. J. Adams. 2020. Shotgun metagenomics reveal a diverse assemblage of protists in a model Antarctic soil ecosystem. Environmental Microbiology 22:4620-4632 and assembly and binning stats from the current work.

Supp. Table 2 – Phenotype Keywords

| Calvin cycle | Antibiotics | Antibiotics (cont.) | Antibiotics (cont.) | AntibioticResistanceGenes | ColdTolerance | Dormancy/persistence |
|---|---|---|---|---|---|---|
| ribulose-1,5-biphosphate carboxylase | Streptomycin biosynthesi | Sulfamonomethoxine | Pseudouridimycin | antibiotic resistance | Trehalose-6-phosphate synthase | SsgA |
| phosphoribulokinase | Tetracycline biosynthesis | Sulfapyridine | 5-azacytidine | antibiotic efflux | otsA | WhiA |
| sedoheptulose bisphosphatase | Vancomycin biosynthesis | Chlortetracycline | Gemcitabine | antibiotic acetyltransferase | trehalose synthase | WhiB |
| ribulose-1,5-biphosphate oxygenase | Erythromycin biosynthesi | Doxycycline | Muramycin | aminoglycoside N-acetyltran | trehalose | WhiD |
| **Arnon-buchanan cycle** | Gentamicin | Oxytetracycline | Liposidomycin | Antibiotic ABC transporter ATP-binding protein | Trehalose-6-phosphate phosphatas | exopolysaccharide production |
| fumarate reductase | Streptomycin | Tetracycline | Capuramycin | msbA | otsB | polyhydroxyalkanoate |
| ferredoxin-dependent 2-oxoglutarate synthase | Trimethoprim | Daptomycin | Tubercidin | MFS transporter | cold shock | polyhydroxyalkanoate synthesis |
| ATP-citrate lyase | Ciprofloxacin | Penicillin | Pyocyanin | RND | CspA | Spo0A |
| **Wood-Ljungdahl pathway** | Difloxacin | Cephalosporin | Tubermycin B | RND efflux pump | CspB | sporulation |
| CO dehydrogenase | Enrofloxacin | Carbapenem | Oxazinomycin | acrB | CspC | **OsmoticStressTolerance** |
| acetyl-CoA synthase | Norfloxacin | Monobactam | Antibiotic Biosynthesis | outer membrane efflux protei | CspD | ProV |
| **Hydroxypropionate (Fuchs-Holo) bi-cycle** | Ofloxacin | Tobramycin | **VirulenceGenes** | tolC | CspE | ProX |
| biotin-dependent acetyl-CoA | Vancomycin | Amikacin | fimbria | beta-lactamase | CspF | BetT |
| propionyl-CoA carboxylase | Lasalocid | Minocycline | pilus | chloramphenicol acetyltransfe | CspG | betaine aldehyde dehydrogenase |
| **3-hydroxypropionate/4-hydroxybutyrate (HP/HB) cycle** | Monensin | Levofloxacin | pili | efflux pump | CspH | BetB |
| 3-hydroxypropionate | Amoxicillin | Moxifloxacin | adhesin | **Motility** | CspI | choline oxidase |
| 4-hydroxybutyrate | Cephapirin | Metronidazole | fimH | flagellum | CspJ | choline dehydrogenase |
| **Dicarboxylate-hydroxybutyrate cycle** | Cefuroxime | Curamycin | inhA | flagellin | β-ketoacyl-ACP synthase II | glycine betaine |
| dicarboxylate-hydroxybutyrate | Penicillin G | Neomycin | biofilm | flagellar assembly | ketoacyl-ACP synthase | **Dessication&RadiationTolerance** |
| **H2_CO_Oxidation** | Clindamycin | Puromycin | biofilm formation | flagellar | ketoacyl-ACP | PolA |
| [NiFe]-hydrogenase | Lincomycin | Amphothericin B | capsular polysaccharid | gliding motility | ribosome-binding factor A | nucleotide excision repair |
| group 1 [NiFe]-hydrogenase | Azithromycin | Nigericin | type III secretion | | polynucleotide phosphorylase | *uvrA* |
| group 2 [NiFe]-hydrogenase | Clarithromycin | Myxin | escV | | Fatty acid desaturase | *uvrB* |
| [FeFe]-hydrogenase | Erythromycin | Rifamycin | type IV secretion | | desA | manganese catalase |
| form I CO dehydrogenase | Tylosin | Neoplanocin | type VI secretion | | desB | rubrerythrin |
| carbon monoxide dehydrogenase | fidaxomicin | Aristeromycin | exotoxin | | desC | dehydrin |
| form II CO dehydrogenase | telithromycin | Toyocamycin | endotoxin | | dhaK | late embryogenesis abundant |
| MMO | Sulfachloropyridazine | Sangivamycin | enterotoxin | | dhaM | late embryogenesis abundant protein |
| sMMO | Sulfadiazine | Showdomycin | neurotoxin | | antifreeze | recA |
| particulate methane monooxygenase | Sulfadimethoxine | Minimycin | toxin | | cold-shock DEAD-box protein A | DNA repair protein RadA |
| MCR | Sulfadoxine | Formycin | quorum-sensing | | ice-binding protein | starvation-inducible DNA-binding protein |
| formate dehydrogenase | Sulfamethoxazole | Pyrazofurin | quorum-quenching | | ice-nucleating protein | Superoxide Dismutase |
| CO oxidase | Sulfamethazine | Malayamycin | | | ice-nucleation | PprA |

Table S2 – Comprehensive list of keywords searched against gene annotation file produced by VEBA, including keywords that recovered 0 hits and were excluded from the main figure (Fig 9). Organized by phenotype category: trace gas metabolism (orange), competitive phenotypes (purple), and extremotolerance

Supp. Table 3 – Metadata

| Site ID | Site Name | Climate Zone (Original) | Samples | % moisture | pH | EC | %Clay | NO3-N (ppm) | Total N (ppm) | Total C (ppm) | C:N Ratio | Total P (ppm) | Elevation (m) | Distance to Coast (km) | Aspect (degrees) |
|---|---|---|---|---|---|---|---|---|---|---|---|---|---|---|---|
| MG | Benson Glacier | Coastal | 3 | 6.64 | 7.3 | 5.70 | 4.52 | 2.99 | 666.7 | 4640.0 | 6.44 | 192.7 | 495.1 | 7.8 | 229 |
| CS | Canada Stream | Coastal | 1 | 12.00 | NA | NA | NA | NA | NA | NA | NA | NA | 54.8 | 11.2 | 317 |
| CN | Cliff Nunatak | Coastal | 2 | 6.37 | 6.7 | 4.08 | 8.54 | 2.53 | 640.0 | 3910.0 | 5.74 | 369.0 | 451.6 | 8.1 | 315 |
| ESLF | F6 | Coastal | 2 | 1.83 | 8.5 | 1.11 | 2.00 | 1.30 | 465.0 | 1630.0 | 3.52 | 958.8 | 17.9 | 5.9 | 265 |
| GV | Garwood Valley | Coastal | 6 | 5.97 | 8.2 | 0.70 | 0.62 | 2.38 | 435.0 | 9323.3 | 25.63 | 616.7 | 625.9 | 10.9 | 215 |
| HH | Hjorth Hill | Coastal | 5 | 5.99 | 7.7 | 1.70 | 1.86 | 2.70 | 534.0 | 4012.0 | 6.87 | 971.2 | 119.0 | 1.0 | 144 |
| Marr | Marr Pond | Coastal | 4 | 4.07 | 7.9 | 13.44 | 3.00 | 19.83 | 505.0 | 4522.5 | 7.68 | 411.0 | 731.6 | 22.9 | 138 |
| MV | Miers Valley | Coastal | 6 | 4.17 | 8.1 | 0.44 | 0.63 | 1.53 | 368.3 | 4506.7 | 12.26 | 558.2 | 214.9 | 9.8 | 272 |
| MS | Mt Suess | Coastal | 3 | 6.93 | 7.7 | 0.32 | 3.02 | 1.30 | 416.7 | 2000.0 | 4.71 | 165.3 | 634.6 | 15.5 | 240 |
| NSLH | Northside Lake Hoare | Coastal | 3 | 1.68 | 8.1 | 2.00 | 4.24 | 1.37 | 193.3 | 2021.7 | 12.11 | 438.0 | 304.1 | 15.4 | 243 |
| WrB | Wright Valley (East) | Coastal | 8 | 2.92 | 8.0 | 10.48 | 3.72 | 50.06 | 391.3 | 1641.3 | 3.88 | 584.1 | 247.8 | 21.7 | 7 |
| MGM | Mount Murray | Inland | 3 | 10.18 | 7.4 | 0.34 | 7.45 | 2.32 | 450.0 | 1500.0 | 3.31 | 236.5 | 767.4 | 11.5 | 324 |
| GR | Mt Gran | Inland | 1 | 3.95 | 7.6 | 1.70 | 0.72 | 3.07 | 300.0 | 1240.0 | 4.13 | 222.3 | 990.7 | 33.6 | 209 |
| TG | Towle Glacier | Inland | 3 | 5.63 | 7.8 | 0.22 | 1.17 | 1.62 | 293.3 | 596.7 | 2.13 | 188.1 | 1015.8 | 42.7 | 119 |
| WrU | Wright Valley (West) | Inland | 8 | 0.70 | 7.9 | 0.28 | 0.73 | 1.31 | 316.3 | 632.5 | 2.01 | 120.7 | 1050.1 | 61.8 | 169 |
| BV | Beacon Valley | Upland | 14 | 0.98 | 7.6 | 4.08 | 1.44 | 83.28 | 347.9 | 774.3 | 2.47 | 345.4 | 2023.3 | 74.8 | 145 |
| UV | University Valley | Upland | 7 | 1.11 | 7.2 | 3.29 | 5.42 | 119.33 | 465.7 | 572.9 | 1.40 | 95.8 | 1689.9 | 72.3 | 211 |
| WV | Wall Valley | Upland | 8 | 0.72 | 7.5 | 2.15 | 2.03 | 25.18 | 320.0 | 615.0 | 1.97 | 218.1 | 1542.6 | 66.0 | 178 |

Table S3 – Averages for environmental variables for all sites. Includes categorical variables for Aridity (based off of % moisture), Elevation, and Distance to Coast (Dist-to-Coast).

**Figure S1 MAG Genome Stats**

A

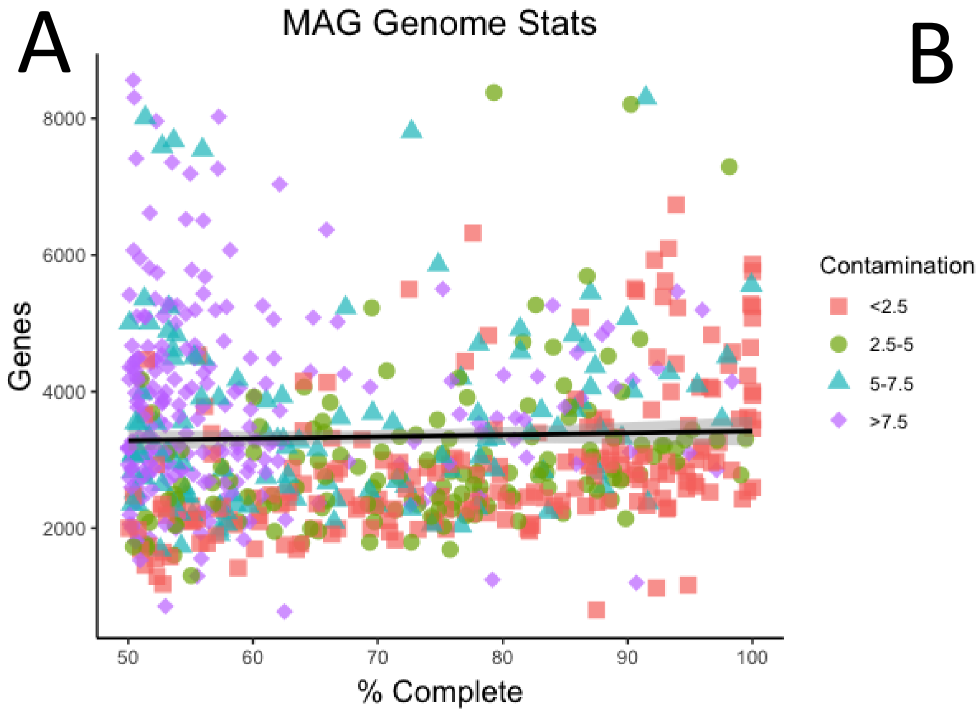


B

**Model - MAGs**

lm(formula = gene_counts ~ Completeness, data = mag_gene_counts)

**Residuals:**

| *Min* | *1Q* | *Median* | *3Q* | *Max* |
|---|---|---|---|---|
| -2584 | -913.6 | -327.5 | 647.3 | 5277.4 |

**Coefficients:**

| | *Estimate* | *Std. Error* | *t-value* | *Pr(>\|t\|)* |
|---|---|---|---|---|
| (Intercept) | 3146.9 | 219.404 | 14.343 | <2e-16 *** |
| Completeness | 2.731 | 3.096 | 0.882 | 0.378 |

**Signif. codes:** 0 '***' 0.001 '**' 0.01 '*' 0.05 '.' 0.1 ' ' 1

**Residual standard error:** 1289 on 702 degrees of freedom
**Multiple R-squared:** 0.001107, **Adjusted R-squared:** -0.0003156
**F-statistic:** 0.7782 on 1 and 702 DF, **p-value:** 0.378

Figure S1 – MAG and SLC genome size (est. gene count). A) Gene counts per MAG by % genome completeness (as estimated by VEBA pipeline software). Each point represents a unique MAG from the dataset. Includes all MAGs with >50% completion and <10% contamination. Point color and shape represents % contamination of MAG assembly. Fitted line shown in black. B) Results of linear model calculation. Adjusted $R^2$ = -0.0003156, p-value = 0.378.

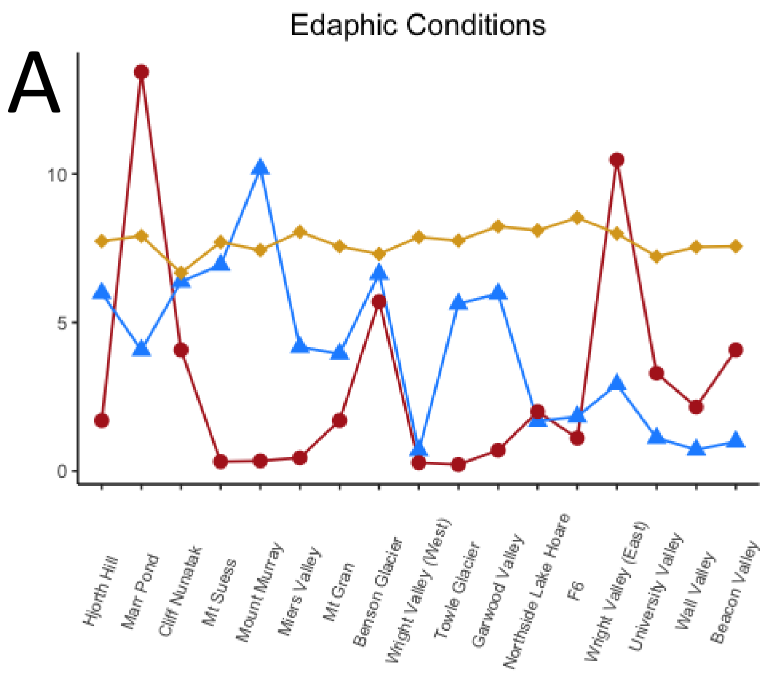


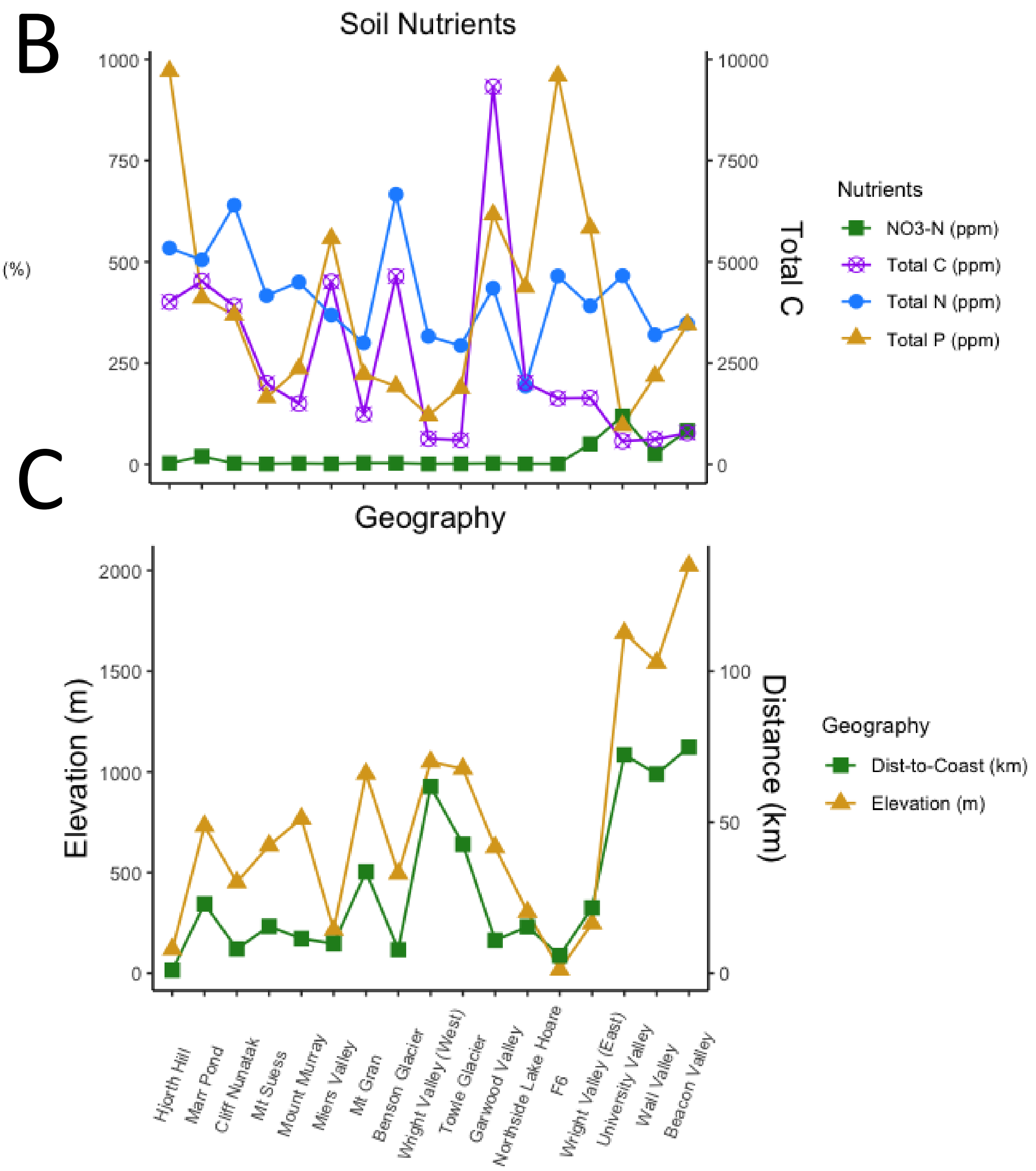


Figure S2 – Metadata overview. X-axis shows site names, y-axis shows values for each parameter. A) Edaphic conditions: EC = electrical conductivity, a proxy for soil salinity. B) Soil nutrients: values for Total C (purple line) are shown on the right y-axis. C) Geography: Dist-to-Coast represents the shortest distance (km) from the sampling site to the Ross Sea coast. Distance to Coast values are shown on the right y-axis.

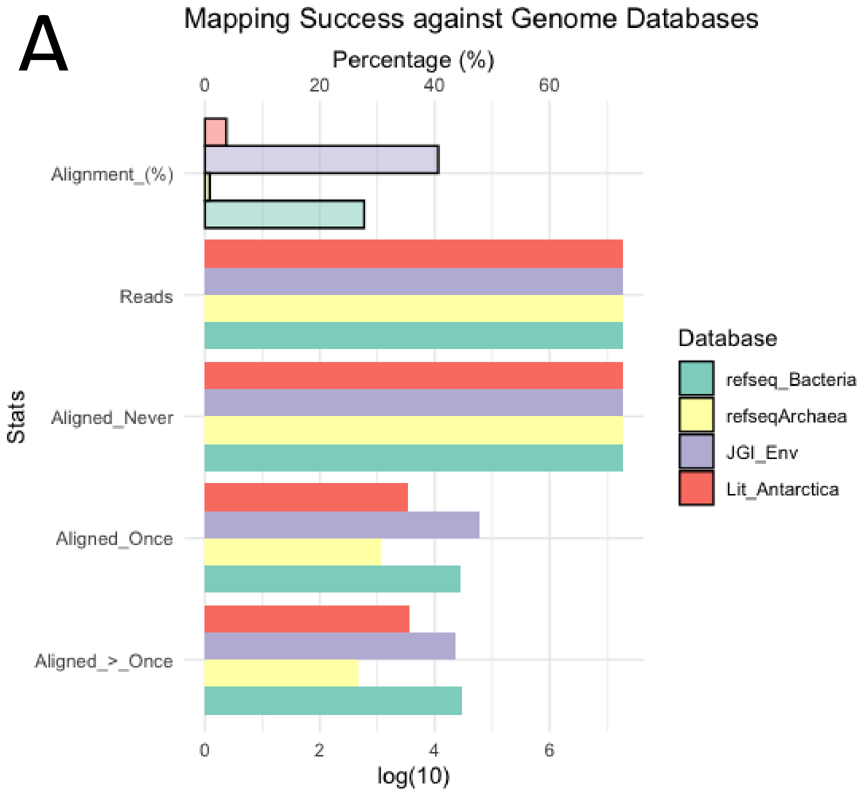


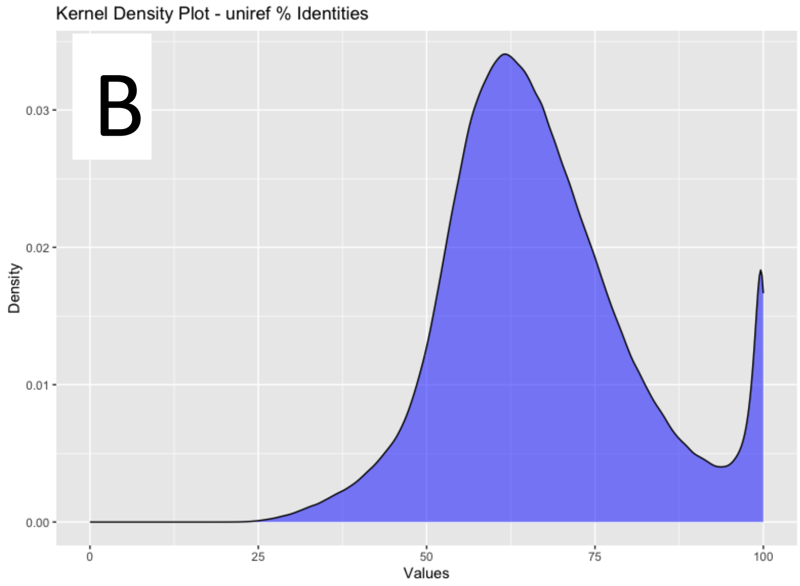


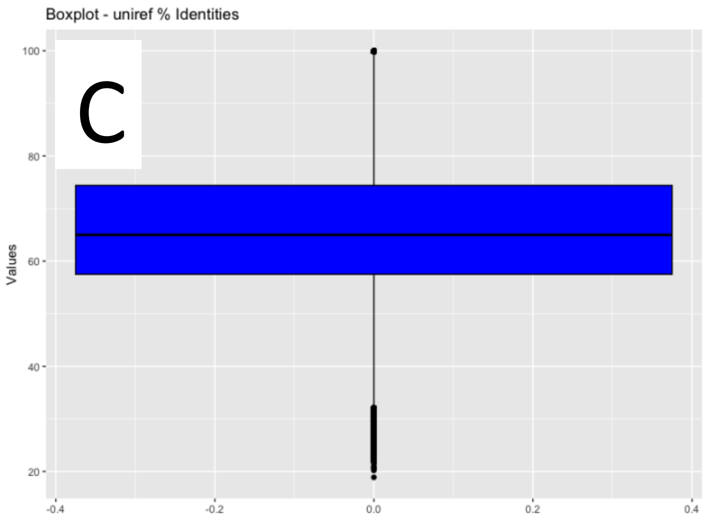


Figure S3 – Alignment Success & Protein divergence. A) Mapping success of metagenomes against four reference databases (2 from NCBI, 2 custom), # total reads, # reads never aligned, # reads aligned once, # reads aligned more than once (scale at bottom). All values besides Alignment % are in $\log_{10}$ scale. BC) Distribution of percent Identity of predicted proteins against UNIREF database – B) Kernal density plot, C) boxplot.

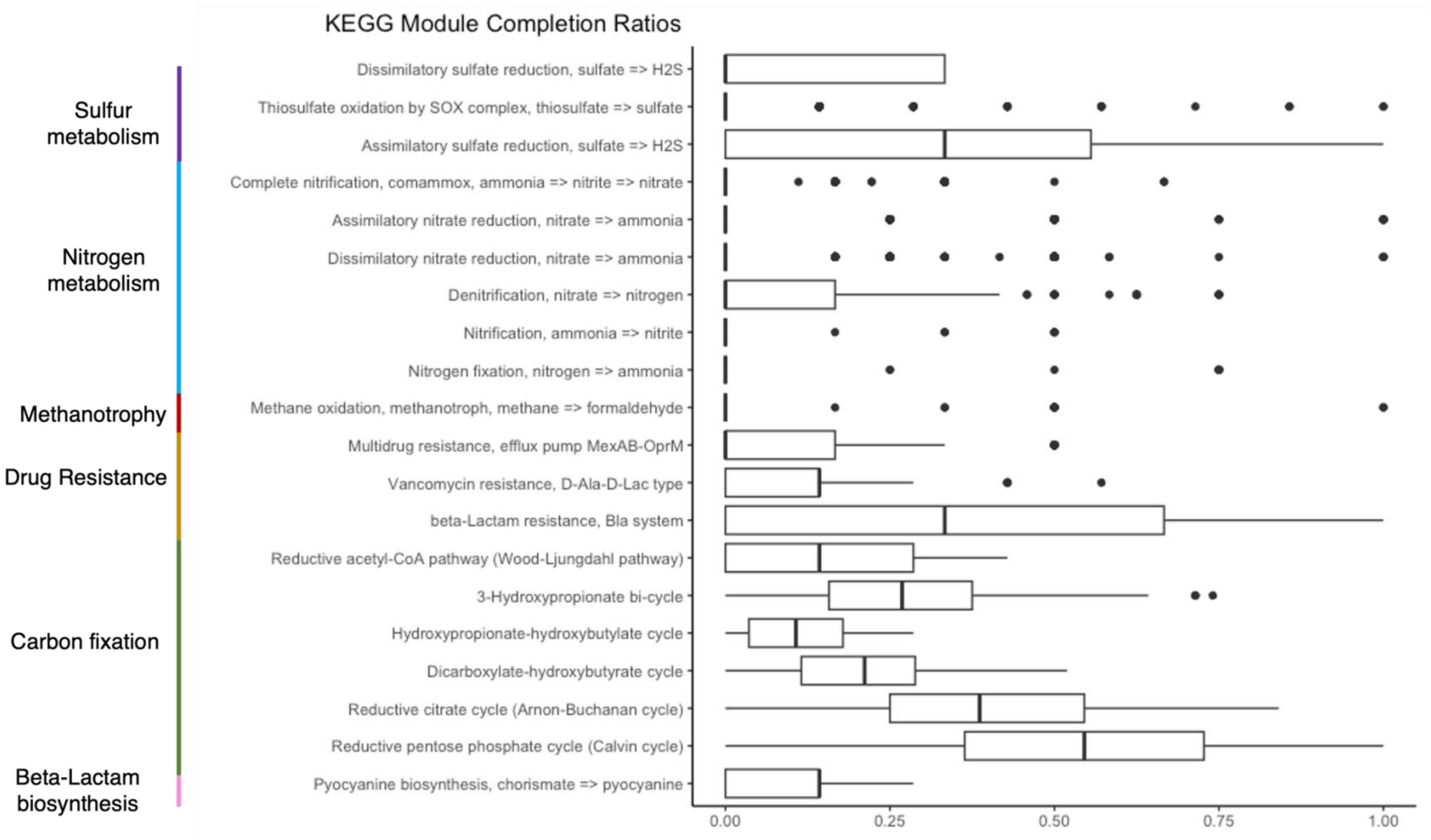


Figure S4 – KEGG Completion Ratio Boxplots by module. Companion plot to Figs 5 & 6 in the main paper: quantitative representation of qualitative visualization of module completion heatmaps in main paper Figs 5 & 6. Only modules of key interest are shown (e.g., modules related to key nutrient metabolism or competition associated modules with non-zero completion). Y-axis shows module names of key interest (near left) and their respective pathway groups (far left). X-axis shows mean % completion of each module.

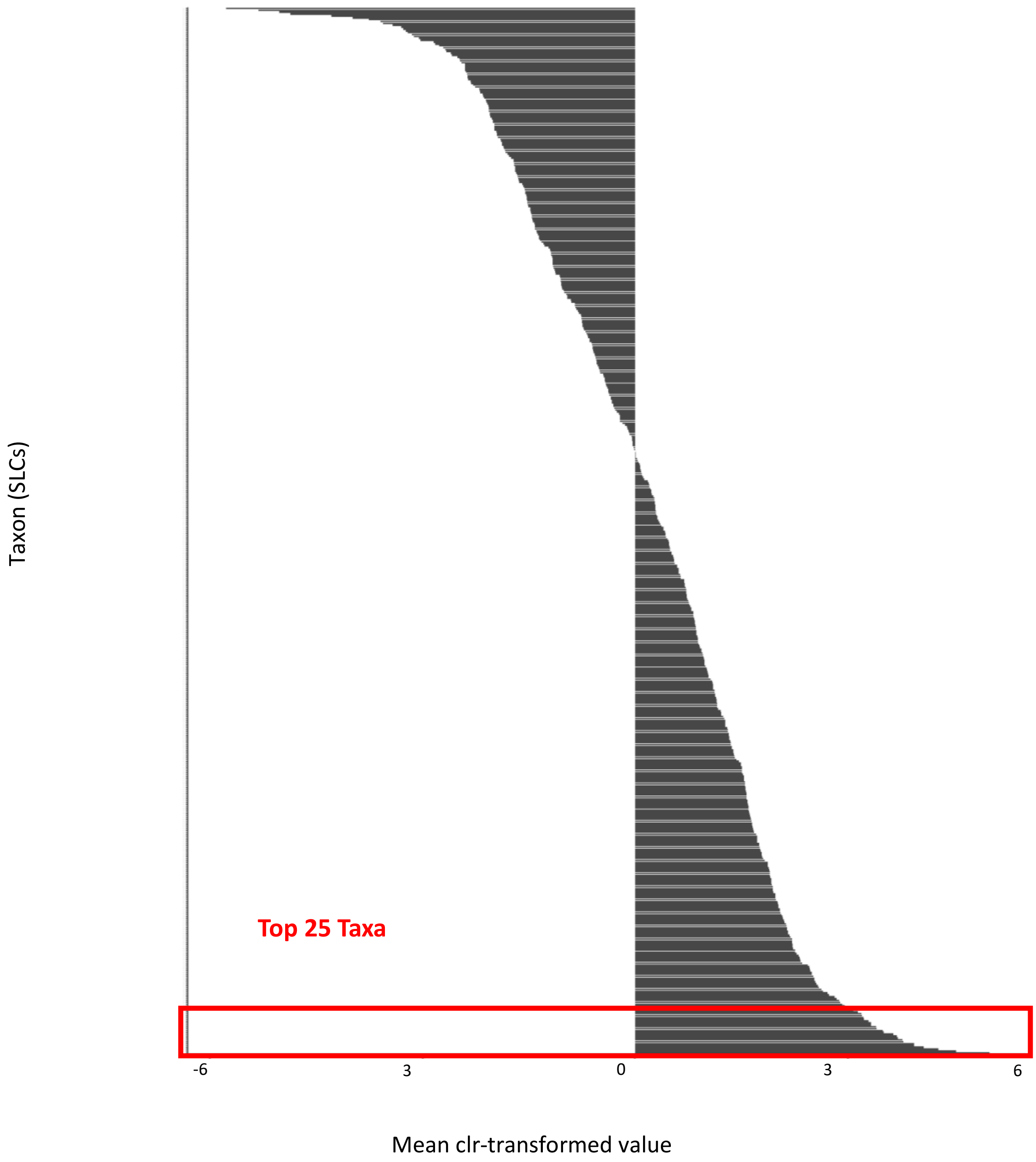


Figure S5 – Taxon relative abundance distribution, with cutoff for Figs5-8. Visualizes all SLCs (486; y-axis) in dataset and their relative abundance (x-axis), after clr-transformation. The least abundant taxa across all sites are near the top, the most abundant are near the bottom. The red box highlights the top 25 taxa for all sites. To determine regional top 25 taxa, only counts from sites assigned to respective regions were averaged, giving a final distribution that was distinct per region (not shown).
